# Matching-Allele-Models explain host specificity and high diversity of Collagen-Like Proteins in a virulent pathogen

**DOI:** 10.1101/2025.03.22.644704

**Authors:** Alix Thivolle, Dieter Ebert

**Affiliations:** Department of Environmental Sciences, Zoology, University of Basel, Basel 4051, Switzerland

**Keywords:** Pasteuria ramosa, host-pathogen interaction, collagen-like protein, genetic basis of infection, genome-wide association study, coevolution

## Abstract

Host specificity is a defining feature of many parasitic interactions, yet the genetic mechanisms behind this specificity remain unclear. *Pasteuria ramosa*, a bacterial pathogen of *Daphnia magna*, provides a model for studying genotype-specific infectivity. We investigate the role of collagen-like proteins (PCLs) in mediating host attachment using genome-wide association studies (GWAS) and presence/absence variation (PAV) analyses. Our results reveal high diversity and lineage-specific patterns within the PCL gene family. For three PCL triplets, we find strong associations with parasite’s attachment phenotypes. Nearly perfect genotype-phenotype matches for certain PCL alleles support the matching allele model (MAM), considered to be an important model of host-parasite coevolution and balancing selection. This is the first molecular evidence of MAM in a pathogen. This study highlights the role of PCL genes in infection specificity, advancing our understanding of host-pathogen coevolution and supporting the Red Queen hypothesis as a driver of genetic diversity.

## Introduction

Antagonistic coevolution between hosts and parasites is recognized as a significant driver of genetic diversity, often maintaining a variety of alleles within host and pathogen populations through mechanisms like negative frequency-dependent selection (NFDS) (Agrawal & Lively, 2002; Ebert & Fields, 2020; Fredericksen et al., 2023; Hamilton, 1980). In host–parasite systems, genetic variation is sustained as parasites continually adapt to common host varieties, resulting in a dynamic cycle where common host resistance alleles and parasite infectivity alleles are selected against in favour of rarer variants. This interaction, often described as Red Queen dynamics, not only drives population differentiation, but also promotes genetic diversity at loci involved in infection and defence (Ebert & Fields, 2020; Woolhouse et al., 2002). Although the study of these interactions has advanced our understanding of evolutionary dynamics, it remains challenging to identify the specific genetic factors that mediate these relationships, particularly in non-model organisms (Nuismer et al., 2017). For an in-depth understanding of how coevolution works, the interacting genes of both host and parasite need to be identified, as the analysis of their variation provides crucial information about the evolutionary mechanisms in place (Dexter et al., 2023; Sironi et al., 2015). Here, we focus on identifying the genes responsible for the phenotypic polymorphisms observed across isolates of a virulent bacterial pathogen.

A model for investigating coevolutionary dynamics is the freshwater crustacean *Daphnia magna* and the obligate bacterial pathogen *Pasteuria ramosa* (Decaestecker et al., 2007; Luijckx et al., 2013), a waterborne disease that transmits horizontally to the susceptible *D. magna* through dormant spores ingested during filter feeding (Ebert, 1996). Genotypes (=clones) of *D. magna* exhibit either complete resistance or complete susceptibility to particular *P. ramosa* isolates, and the infection’s outcome is governed by highly specific genotype-by-genotype interactions (Carius et al., 2001; Luijckx et al., 2011). In *P. ramosa*, infectivity is likely mediated by a diverse family of collagen-like proteins (CLPs) that have been widely implicated in host–pathogen interactions in other bacteria such as *Streptococcus pyogenes, Bacillus anthracis*, and *Legionella pneumophila* (Bachert et al., 2015; Lukomski et al., 2000, 2017; Sylvestre et al., 2002; Vandersmissen et al., 2010). These proteins are characterized by the Gly-Xaa-Yaa repeat structure, essential for forming the stable triple-helices that confer strength to extracellular matrices (Frantz et al., 2010; Qiu et al., 2021). Collagen-like proteins have been linked to various bacterial functions, including biofilm formation, immune evasion, and host cell adhesion, often acting as adhesins for pathogens to attach to host tissues and initiate infection (Bachert et al., 2015; Oliver-Kozup et al., 2011; Pizarro-Guajardo et al., 2014; Reid, 1993). The PCL gene family is highly expanded in *P. ramosa*, with many PCLs arranged in triplets that may play an important role in pathogen–host interactions (Andras et al., 2020; Dexter et al., 2023; Huessy et al., 2023; McElroy et al., 2011). One specific PCL gene, PCL7, was found by a recent study to contain polymorphisms that correlate with the bacterial spores’ attachment to *D. magna* (Andras et al., 2020; Huessy et al., 2023). It remains unclear, however, why not all bacteria have this attachment-allowing genotype. The authors speculate that bacterial genotypes that are unable to attach to one host may be able to attach to other host genotypes, such that various *P. ramosa* genotypes are specialized to attach to specific host genotypes, as suggested in gene-for-gene and matching-allele models of host–parasite interaction (Agrawal & Lively, 2002; Luijckx et al., 2013).

In this study, we investigate the diversity and distribution of PCL genes in *P. ramosa* and explore their association with attachment phenotypes across different *D. magna* genotypes. Using a combination of genome-wide association studies (GWAS) and gene presence/absence variation (PAV) analysis, we examine a geographically diverse collection of *P. ramosa* isolates, finding substantial variation in the PCL gene family, including the organization of PCLs into lineage-specific clusters and significant differences in nucleotide diversity among PCL triplets. We identify PCLs specifically linked with attachment to certain host tissues---such as the foregut and external abdomen--and observe variability in attachment specificity depending on the *P. ramosa* lineage. These results suggest that lineage-based differences in PCL gene content and sequence diversity may be critical for facilitating coevolutionary interactions between *D. magna* and *P. ramosa*. Our findings coincide with a matching-allele-model, giving strong support for the Red Queen model of antagonistic coevolution.

## Results

### PCL Diversity

PCLs are known to play a role as interactors between *Pasteuria ramosa* and *Daphnia magna*. According to previous research, they generally fall into five groups: four groups of PCLs form tight genetic clusters, while the remaining are unclustered (McElroy et al., 2011; Thivolle et al., 2024). PCL groups 1, 2, and 3 always occur in triplets in the genome, each containing one PCL from each group 1, 2, and 3, respectively. These triplets always occur in the same order and orientation within each triplet. So far, 13 different triplets have been detected and are hypothesized to be involved in *P. ramosa* spore attachment (Andras et al., 2020; Dexter et al., 2023; Huessy et al., 2023). The role of PCL group 4 and the unclustered PCLs is still unknown. Within *P. ramosa* genomes, PCLs genes and, in particular, PCL triplets are diversity hotspots (Figure 1A), as is consistent with balancing selection (Charlesworth, 2006; Ebert & Fields, 2020; Nielsen, 2001). The nucleotide diversity per PCL is highly variable, with strong differences among the five PCL groups (Figure 1B, C). Group 4 PCLs are highly conserved with a low nucleotide diversity (mean: 0.001617, sd: 0.0007268), very similar to the genomic background (0.00158). The three PCLs groups found in triplets have high diversity and show little difference between them. The ungrouped PCLs fall in between the estimates for the triplets and group 4. There is also high variation in the diversity between triplets (Figure 1D): some are highly conserved as PCL03-04-05, PCL52-53-54 with pi 0.000476 and 0.000637; others are nearly 100 times more diverse, such as PCL36-37-38 and PCL26-25-24 (pi= 0.037545, pi= 0.032437). This diversity is also reflected in the number of alleles observed in our sample per PCL (Figure 1C and 1E). A discrepancy appeared between the low number of alleles in PCL 12-13-14 and its high nucleotide diversity; however, closer examination showed that this triplet is similar to another triplet (Pcl36-37-38), causing read mis-mapping and false SNP calls. PCL diversity is also visible in terms of gene presence or absence. Figure 1F illustrates how the distribution of PCL presence across isolates is highly triplet-specific: about half (6 of 13) of all triplets are found in almost all isolates, while the others occur in a range of 15 to 80 % of isolates. There was no consistent relationship between the conservation of a PCL and its prevalence. For example, while PCL03-04-05 is highly conserved and present in most isolates, the equally-conserved PCL52-53-54, is found in less than a quarter of isolates.

**Figure 1:**
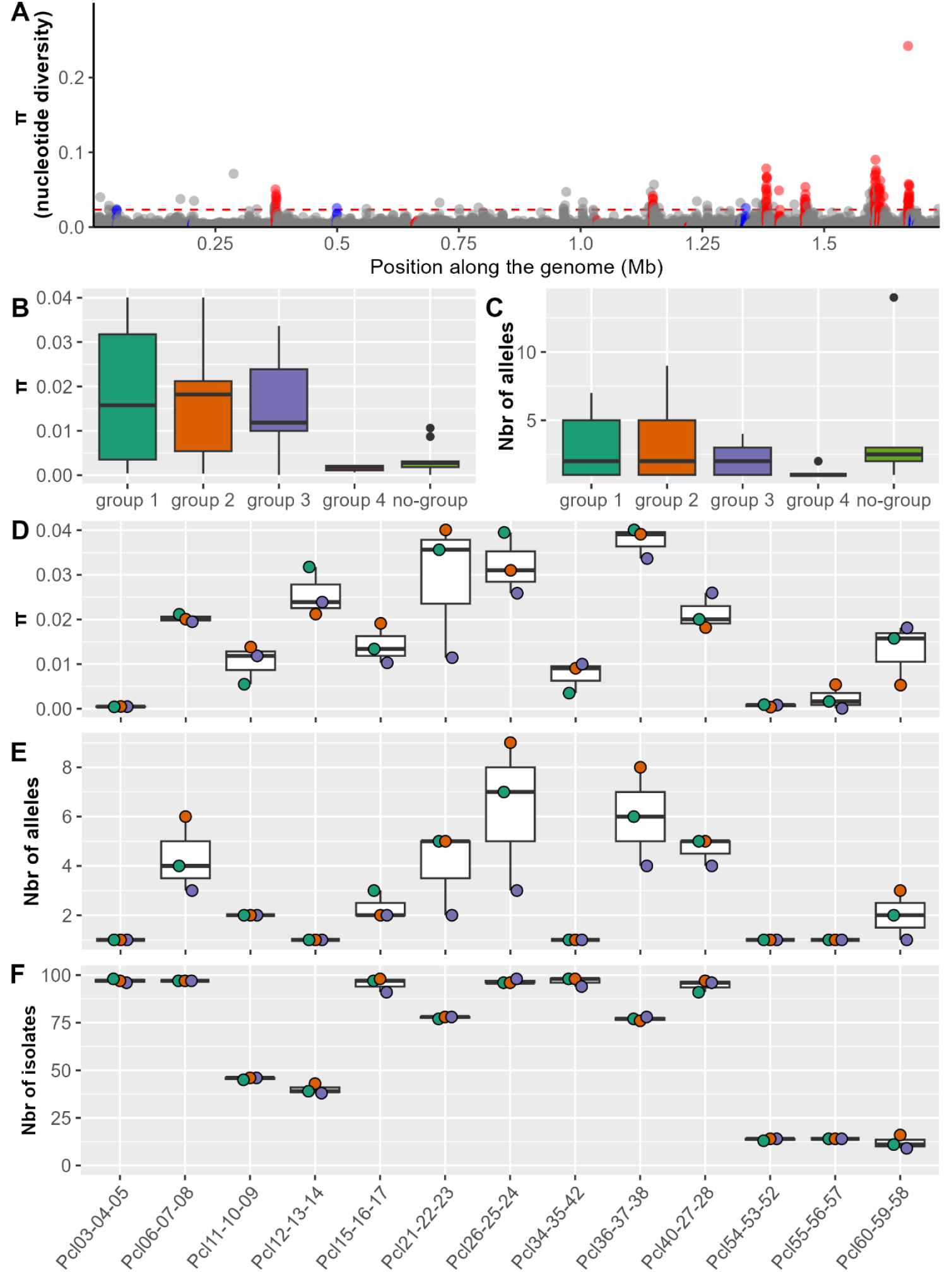
Highest diversity is observed in PCL regions, though not all PCLs exhibit high diversity. **A**. Nucleotide diversity across the *P. ramosa* genome, calculated using a sliding window of 100 bp. The dashed red line represents the 99th quantile, and the solid line marks the 95th quantile. Red dots indicate positions of nucleotides belonging to a PCL-triplet (PCL cluster from groups 1, 2 and 3 (McElroy et al., 2011)); blue dots represent other PCLs. **B**. Box plot showing nucleotide diversity for each of the five PCL groups: the four known clusters and the fifth for ungrouped PCLs. **C**. Similar to Panel B, this box plot displays the number of alleles per PCL per group. **D and E**. Panels showing nucleotide diversity and number of alleles for each PCL triplet. Dots represent the three PCLs within each triplet, with color coding denoting group affiliation consistent with panels B and C. **F**. Panel showing the number of isolates that exhibit at least one copy of a PCL. Nucleotide diversity was calculated using the GWAS dataset of 91 isolates. Allele numbers were detected using isolates that met our criteria of assembly completeness and multi-infections, leading the number to 98 isolates (Supplementary File 1). The same isolates were used for panel F.

### PCL and Lineages

The SNP-based phylogenetic tree confirms the lineage clustering of the isolates as previously described (Dexter et al., 2023), except that a fourth lineage appeared as more diverse *P. ramosa* were added; we named it delta (Figure 2 left). Also, the absence of certain PCLs triplets appeared to be largely, but not exclusively, lineage-specific (grey in Figure 2). For instance, PCL triplets PCL11-10-09 and PCL14-13-12 were primarily found in the beta lineage, while PCL54-53-52/55-56-57 and PCL60-59-58 were predominantly associated with the alpha lineage. PCL21-22-23 is generally absent in the alpha lineage. These patterns reveal several positive associations between PCL triplets, but the presence/absence patterns of triplets can also be negatively correlated. For example, triplets PCL60-59-58 and PCL21-22-23 are never found together within the same genome. This may be due to genomic positioning, as a copy of PCL60-59-58 in isolate P54 occupies the same location as PCL21-22-23 in the genome of isolate C1.

**Figure 2:**
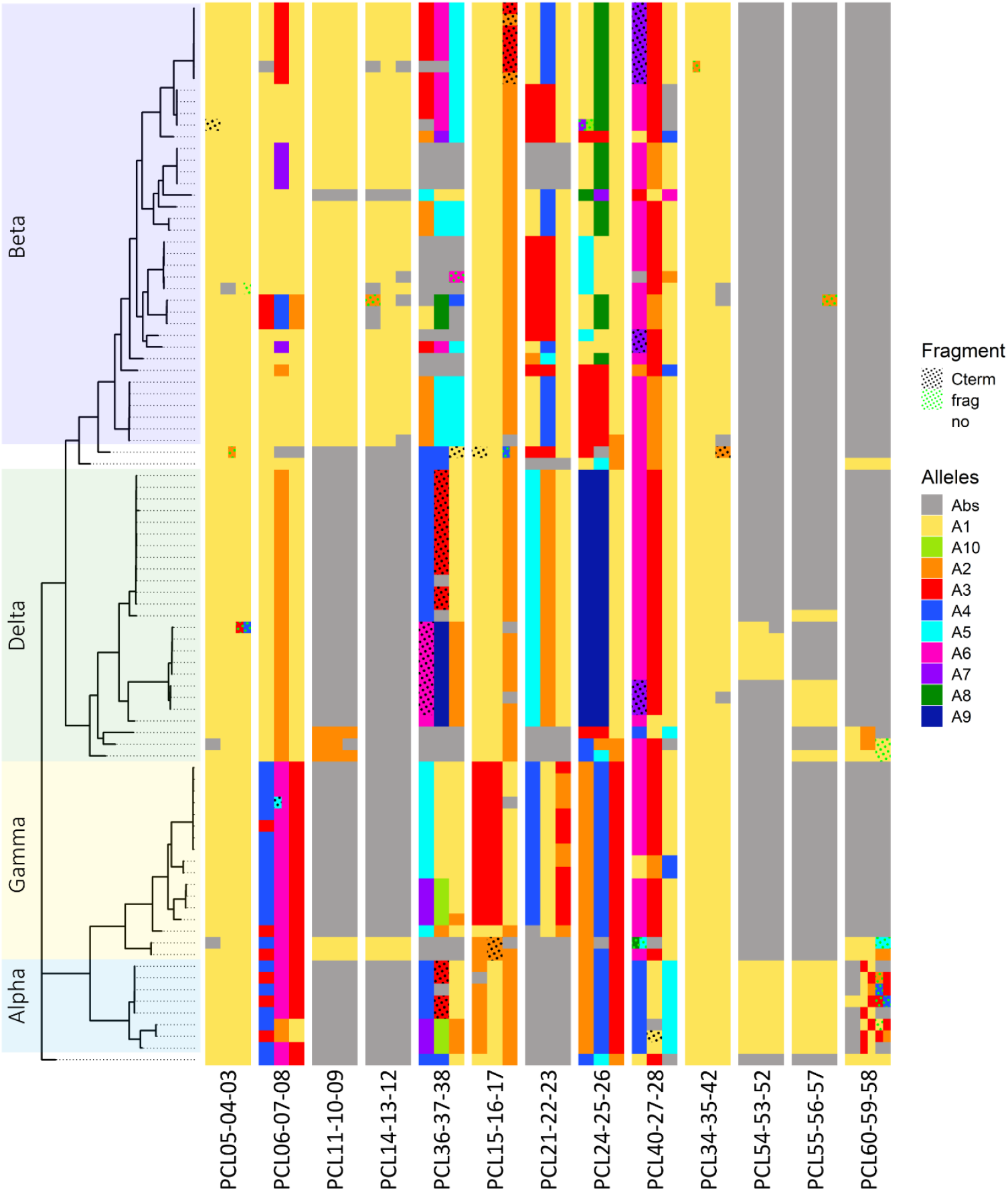
Phylogenetic structure and PCL alleles. **Left Panel:** Unrooted phylogenetic tree based on SNPs data, highlighting four lineages—alpha, beta, gamma and delta—and omitting isolates that did not classify easily into a lineage. **Right Panel:** Matrix depicting the presence, absence (in grey), and different alleles for each PCL organized by PCL triplets. PCL triplets are composed of one PCL each from group 1, 2 and 3, oriented in the same direction. PCLs can have up to 10 different alleles, defined here as PCL sequences with a homology over 95%. Half boxes indicate PCLs that are duplications; green dots show PCL fragments; black dots indicate PCL fragments that lack only the C-terminal part of the protein.

The allelic diversity of PCL, like presence/absence patterns, also revealed lineage specific patterns (colors in Figure 2). Some PCL alleles are unique to certain lineages or sub-lineages, while others are shared across lineages. For example, allele 2 of PCL07, is mainly found in delta, and allele 1 of PCL16 is found in both the beta sub-lineage and alpha lineage, whereas the gamma lineage displays a different allele.

### Spore adhesions assay and phylogenetic tree

Fluorescently labelled *P. ramosa* spores were tested for tissue tropism to determine the host sites to which they attach. These sites show considerable variability across isolates (**Error! Reference source not found**.), both across and within lineages (Figure 3). For instance, isolates in the beta lineage exhibited a pronounced tendency to attach to the host foregut, with all 41 of its isolates attaching to the foregut of at least one host genotype and most (34 out of 41) with attachment proportions above 80% (mean = 0.898, sd = 0.104) (blue in Figure 3). Still, this attachment was highly specific: no *P. ramosa* isolate attached to the foregut of every host genotype, and only a few overlapped across hosts (of 9 host genotypes, 2 isolates attached to the foreguts of 2, 3 to 3, 11 to 4, 14 to 5, 8 to 6, and 3 to 8 hosts). Additionally, 15 beta isolates exhibited secondary attachment to other sites but at lower proportions or limited to a single host clone.

**Figure 3:**
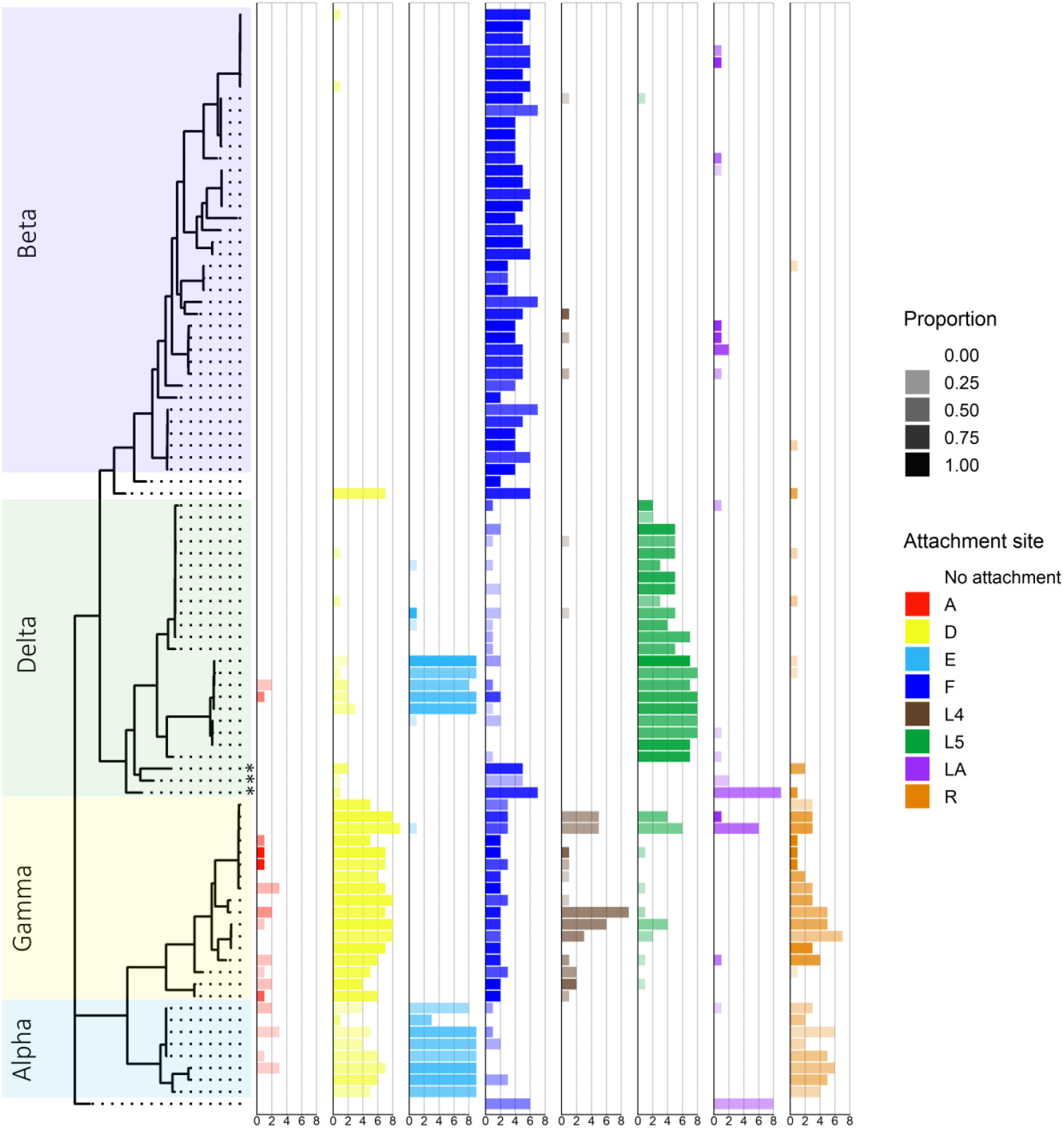
Lineage-specific tropism for attachment sites, ignoring host clone specificity. **Left Panel:** Phylogenetic tree as in Fig. 2. **Right Panel:** Bar plot representing attachment for each isolate to specific host attachment sites. Attachment sites are color-coded, with color shading indicating the proportion of observed attachments. Bars ranging from 0 to 9 indicate the number of hosts the isolate is able to attach to at this site.

In the delta sub-lineage, there was a marked preference to attach to the trunk limb 5 (L5 in green in Figure 3), with 24 of 28 isolates attaching to at least one host at L5. However, attachment strength was lower, with only nine of 24 isolates showing proportions above 80% (mean = 0.732, sd = 0.142). Attachment in delta was also less specific, with more isolates binding to multiple host clones (of 9 hosts, 2 isolates attached to 2, 2 to 3, 1 to 4, 7 to 5, 5 to 7, and 7 to 8 hosts). Only four delta isolates did not attach to any other host site, while six also attached strongly to both the external abdomen and the distal hindgut of some host clones. Three isolates demonstrated a divergent infectotype, attaching to sites other than L5 (Figure 3 the three isolates marked with an asterisk). Notably, these isolates also displayed mixed ancestry in the admixture plot (**Error! Reference source not found**., the three isolates marked with an asterisk).

Gamma lineage isolates were distinguished by robust but non-host-specific attachment to the distal hindgut (D in yellow in Figure 3) across most host clones: 19 isolates attached to the hindgut of at least one host genotype, with nine displaying attachment proportions over 80% (mean = 0.789, sd = 0.069). Attachment specificity was also evident, with gamma isolates showing strong attachment to the foregut of two of the nine host clones. Some gamma isolates were also the only isolates to attach to trunk limb 4 (L4 in dark brown in Figure 3).

The alpha lineage was characterized by strong but non-host-specific attachment to the external abdomen across multiple host clones: nine isolates attached to at least one host, although only two isolates exceeded 80% attachment (mean = 0.647, sd = 0.134). A subgroup of beta attached to the same site. Additionally, isolates in the alpha lineage exhibited weak, non-specific attachment to the distal hindgut (D) and rectum (R).

### Genome-wide association studies based on SNPs

We initially identified 14,195 genetic variants, which were subsequently filtered using several stringent criteria, resulting in 6,149 single nucleotide polymorphisms (SNPs) for downstream analyses. Population structure analysis revealed a distinct clustering pattern corresponding with genetic lineages (**Error! Reference source not found**.), as has been seen in earlier studies (Dexter et al., 2023). Given this clear, lineage-based clustering, a correction was applied categorizing individuals into four groups (**Error! Reference source not found**.), to account for lineage-related confounding factors during the genome-wide association studies (GWAS).

We focused our GWAS on describing the foregut attachment signal, as it was the most prominent and interpretable result in our dataset. Signals associated with other attachment sites were weaker and presented greater interpretive challenges. We first aimed, and succeeded, in replicating a previously reported association for foregut attachment in the HU-HO-2 host (Figure 4) (Andras et al., 2020), detecting a very strong signal in the PCL triplet PCL06-07-08, with the most pronounced effect localized in PCL07. Allele analysis revealed that alleles 1 and 7 were associated with this attachment phenotype, with seven specific non-synonymous SNPs identified (Table 1). Similar results were found for host genotype RU-HA-1, a genotype isolated several thousand kilometers east of the Hungarian clone HU-HO-2.

**Figure 4:**
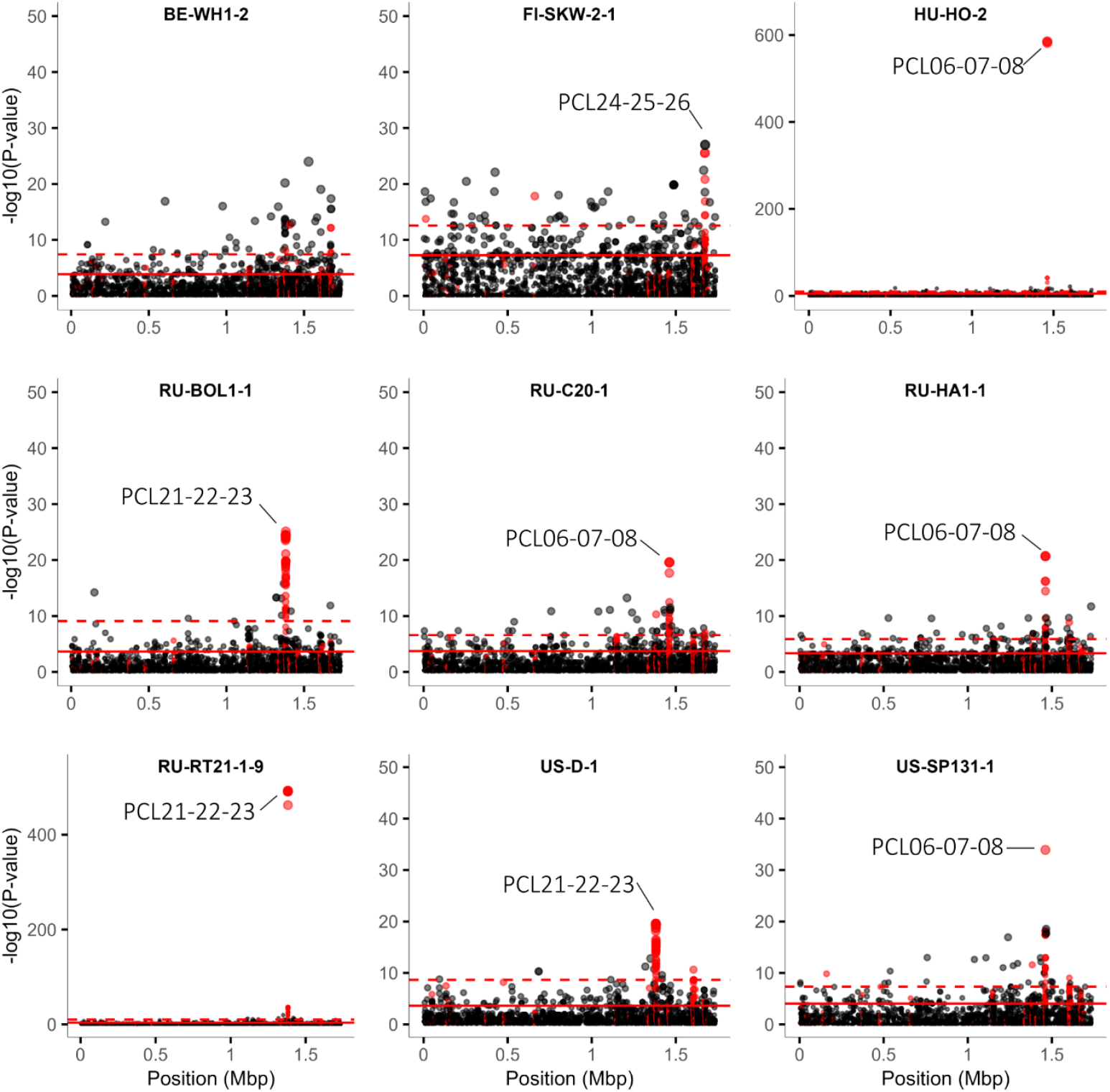
GWAS analysis of foregut attachment across nine host clones. Each panel displays a Manhattan plot of GWAS results for a specific host clone. The dashed line marks the 99th quantile, while the solid line indicates the 95th quantile. Red dots highlight SNPs within PCL-coding regions.

**Table 1:**
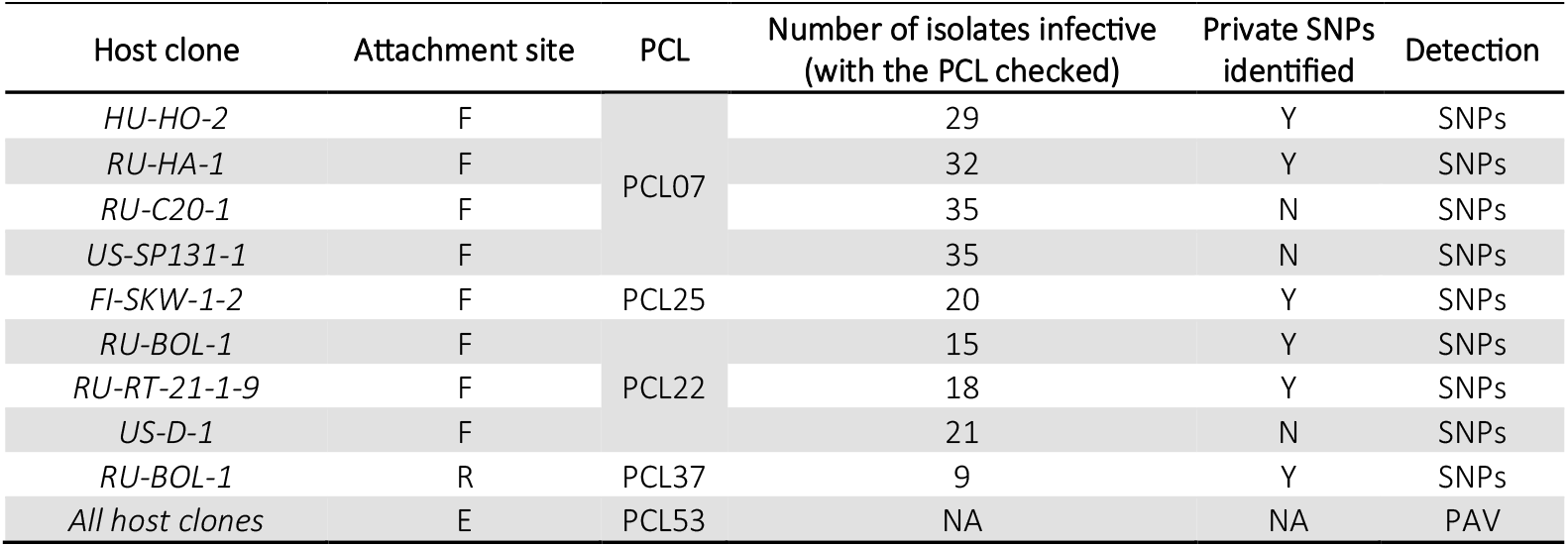
Summary results.

| Host clone | Attachment site | PCL | Number of isolates infective<br>(with the PCL checked) | Private SNPs<br>identified | Detection |
| --- | --- | --- | --- | --- | --- |
| HU-HO-2 | F | PCL07 | 29 | Y | SNPs |
| RU-HA-1 | F |  | 32 | Y | SNPs |
| RU-C20-1 | F |  | 35 | N | SNPs |
| US-SP131-1 | F |  | 35 | N | SNPs |
| FI-SKW-1-2 | F | PCL25 | 20 | Y | SNPs |
| RU-BOL-1 | F | PCL22 | 15 | Y | SNPs |
| RU-RT-21-1-9 | F |  | 18 | Y | SNPs |
| US-D-1 | F |  | 21 | N | SNPs |
| RU-BOL-1 | R | PCL37 | 9 | Y | SNPs |
| All host clones | E | PCL53 | NA | NA | PAV |

Despite a strong signal in PCL07 for attachment to host clones RU-C20-1 and US-SP-131-1, it was challenging to link this signal with specific genotypes or alleles. By restricting the GWAS to beta lineage *P. ramosa*, we saw clear correlations between the attachment phenotype and alleles 3, 4, and 7 of PCL07. However, these alleles contained no private SNPs, making it difficult to identify the specific genetic variants responsible for this phenotype (see Table 2A for PCL07 attachment summary). Despite the fact that the segregation of alleles according to phenotype is imperfect, the substitution of one allele with another clearly inverts the infection profile across host genotypes, making it possible to identify the pattern of a matching-allele model.

**Table 2:** Matching allele model. Infection matrix highlighting the segregation of the phenotype-allele interactions. The matrix shows the outcomes of the interaction between host genotypes (rows) and parasite genotypes (columns). The A panel depict the role of PCL07 while the B panel is focusing on the role of PCL22 The value in the box gives the percentage of isolates presenting the PCL allele (column) able to attach to site specific host (row). Boxes with a black background are showing a strong pattern of specific attachment, white are non-attachment, grey is mix. In the panel A the allele A5 is not mentioned because it appears to be only a fragment present in only one isolate.

| A |  | Parasite PCL07 Alleles |  |  |  |  |
| --- | --- | --- | --- | --- | --- | --- |
|  |  | A1 | A2 | A3/A4 | A6 | A7 |
| Host | HU-HO-2 | 100 | 0 | 0 | 0 | 100 |
|  | RU-HA-1 | 100 | 6.5 | 10 | 0 | 100 |
|  | RU-C20-1 | 4.3 | 6.5 | 100 | 65.4 | 100 |
|  | US-SP131-1-1 | 4.3 | 6.5 | 100 | 65.4 | 100 |

**Table 2 : Matching allele model**
| B |  | Parasite PCL22 Alleles |  |  |
| --- | --- | --- | --- | --- |
|  |  | A3 | A4 | Other Alleles |
| Host | RU-BOL-1 | 88.2 | 0 | 0 |
|  | RU-RT21-1-19 | 0 | 100 | 0 |
|  | US-D-1 | 16.6 | 100 | 0 |

A strong connection was found between the PCL21-22-23 triplet and attachment to host clones RU-BOL1-1 and RU-RT21-1-9 (Figure 4), which exhibited nearly perfect mutual exclusivity in their association patterns. One PCL22 allele was linked to attachment to RU-BOL1-1 (PCL22_A3), while a second allele was associated with attachment to RU-RT21-1-9 (PCL22_A4, Table 1, Table 2B). Each allele contained distinct private SNPs, mainly in the C-terminal domain, that resulted in non-synonymous amino acid changes (Table 1) and suggest functional differentiation between the alleles for host attachment. Table 2B shows a nearly perfect segregation of the allele according to the phenotype of attachment, leading to a very strong genotype matching model.

Similarly, we identified a strong signal within the PCL21-22-23 triplet associated with attachment to the foregut of host clone US-D-1; however, pinpointing a specific genotype responsible for this attachment proved challenging. No common or private SNPs were identified to explain this attachment specificity.

A weaker association exists between the 8th allele of PCL25 and attachment to the foregut of host clone FI-SKW-2-1 (PCL25, Table 1, Figure 4).

### GWAS based on presence/absence variation

We conducted an alternative GWAS based on gene presence/absence variation (PAV) using PPanGGOLiN followed by Scoary. PPanGGOLiN constructs pangenomes and clusters genes into orthogroups based on sequence homology and genomic neighborhood, allowing some PCL alleles (e.g., PCL07) to cluster in a single gene family, while others (e.g., PCL22) split across multiple gene groups. We applied Scoary2 to the presence-absence matrix generated by PPanGGolin and the infectotype matrix previously binarized with a cutoff at 50 % to connect genes with traits,

Through Scoary, we found a strong and consistent correlation between the presence of PCL53 and attachment to the external abdomen across host clones, though association strength varied by host (**Error! Reference source not found**.). The second and third hits across host clones were consistently PCL52 and PCL54, with the fourth hit located near this triplet.

Our GWAS approach resulted in several hits to triplet PCLs, but none to any non-triplet PCL (group 4 and ungrouped), nor to any other gene. This coincides with previous studies. On the other hand, five of 13 PCL triplets were found, at least in one case, to be associated with an attachment phenotype (Table 1). A larger or a differently structured sample, or a sample collected at a different time point, may reveal a role for more PCLs in this system.

## Discussion

To understand the process of coevolution, it is necessary to pin-point the underlying genes in both antagonists. Here we aimed to find the genes in the bacterial parasite *Pasteuria ramosa* responsible for attaching to its host, the planktonic crustacean *Daphnia magna*, with which it is known to coevolve. We hypothesized that collagen-like genes are the key players for host attachment and sequenced more than 100 genomes of *P. ramosa* to analyze genomic and phenotypic data. Besides this species’ well-known phenotypic variation, we observed ample genetic variation among isolates, as measured by both nucleotide diversity and gene presence/absence polymorphisms. While several PCL gene triplets stood out as hot-spots of nucleotide diversity and allele richness, other triplets did not, but showed presence/absence polymorphisms. Both these types of genetic variation showed strong associations with attachment phenotypes, but only in connection with PCL triplets. The main phylogenetic lineages showed support for genetic recombination and evidence of matching-allele models of infection specificity. These results are in support of the Red Queen model of coevolution.

### Lineages and Phenotypic Characterization

Our study confirms previous findings from a single population of *Pasteuria ramos*a and its three main phylogenetic lineages (Dexter et al., 2023) and extends it using a sample collected across many populations and discovering a fourth lineage. We also provide strong evidence for the presence of lineage differences in the phenotypic data from these isolates. Specifically, we observe that the beta lineage attaches predominantly to the foregut, while delta shows a marked preference for the trunk limb (L5) attachment. In contrast, the gamma and alpha lineages exhibit a more versatile attachment profile, with gamma lineage isolates targeting the foregut or distal hindgut, and alpha lineage isolates attaching predominantly to the external abdomen or distal hindgut. Despite these specific attachment profiles, our data suggests that recombination in the *P. ramosa* genome likely facilitates the shuffling and exchange of genes related to attachment and enhances adaptability in response to host diversity.

Recombination and horizontal gene transfer (HGT) in pathogenic species generates diversity, as observed in *Plasmodium falciparum*, where recombination facilitates the fusion of *var* genes, expanding the pool of antigenic genes used for immune evasion (Kirkman & Deitsch, 2014). Similarly, in *Borrelia burgdorferi*, the bacterium responsible for Lyme disease, recombination in the genes’ encoding surface antigens enhances its ability to adapt to a wide range of host species (Akther et al., 2024). Although we did not find mosaic genes in the PCLs or clear signs of recombination in our *P. ramosa* sample genomes, we did observe a strong decay of LD across about 5 kb (**Error! Reference source not found**.), consistent with past recombination.

Diversity can also be created by uptake of genes from the environment (Arnold et al., 2022; Chen et al., 2014; Dutta & Pan, 2002), as we see in *Helicobacter pylori*, which uptakes environmental DNA and incorporates it into its genome. In this bacterium, recombination is common and plays a significant role in shaping the genomes of co-infecting strains (Jackson et al., 2020; Suerbaum & Ailloud, 2023; Woods et al., 2020). Recombination and HGT are thought to shuffle gene repertoires and facilitate gene exchange within and among lineages in our *P. ramosa* sample, as well. For instance, triplets PCL52-53-54 and PCL55-56-57 appear in different combinations in various lineages and may be mobilized through recombination or horizontal gene transfer during naturally occurring co-infection. Alternately, it is possible that these triplets were present in the universal ancestor whose genes were lost in different branches but retained by the alpha lineages and a sublineage of delta. A similar case may exist for triplet PCL60-59-58: it cannot be easily ruled out that this triplet is ancestral and was lost in various branches of the tree.

### Red queen model and Genotype matching model

The remarkable functional diversity observed among the sampled isolates underscores the dynamic nature of host–parasite coevolution. This diversity, characterized by distinct infectotypes and nearly perfect exclusivity in some genotype–phenotype associations, is a prerequisite for the form of negative frequency-dependent selection that underpins the Red Queen model of coevolution. According to this model, antagonistic coevolution between hosts and their parasites maintains genetic variation and explains the persistence of sexual reproduction despite its costs (Lively, 2010). The necessary genetic variation is created either through continuous genetic recombination and sexual reproduction or from a large pool of standing genetic variation (and possibly rare genetic recombination). For host–parasite coevolution to drive such patterns of diversity, specificity in interactions is crucial. Parasites must be adapted to infect particular host genotypes, while hosts must resist specific parasite genotypes. The matching-allele model (MAM) provides a framework for this specificity, positing that resistance or susceptibility can be determined by allelic variation at a single locus (Agrawal & Lively, 2002).

In our study, the observed patterns of genotype–phenotype matching closely align with the MAM framework. For example, the allele PCL22_A3 was associated exclusively with attachment to the foregut of a specific host lineage, while PCL22_A4 exhibited a distinct and exclusive attachment phenotype to a different host lineage. Other PCL22 alleles showed no evidence of foregut attachment, but may do so in other hosts that were not part of our test panel. This pattern—where replacing one allele with another can invert the infection profile across host genotypes—is a hallmark of the genotype-matching model and echoes similar findings for the host in the *Daphnia magna–Pasteuria ramosa* system (Luijckx et al., 2013). A MAM was also observed for PCL07, answering the question of what would prevent previously discovered infective haplotypes from replacing non-infective haplotypes (Andras et al 2020). However, a new paradox seems to have emerged, in that allele A7 was able to infect all host genotypes, while allele A2 was not infectious for any. We speculate that the host genotypes resistant to A7 and susceptible to A2 were not included in our small panel of nine host genotypes.

This interplay of host–parasite diversity is at the very center of the Red Queen model, where genetic variation is both a driver and a consequence of the adaptive dynamics. The coupling of host resistotypes with parasite infectotypes in such systems illustrates the intricate and reciprocal nature of coevolution, while the specificity at the genotype level helps shape evolutionary trajectories on both sides of the interaction.

The presence/absence polymorphisms discovered here for seven of the 13 PCL triplets presents a different form of genetic diversity. One triplet, PCL54-53-52, exists in a single allele that is shared across different lineages: although it is predominantly found in the alpha lineage, it is also detectable in other lineages such as a sub-clade of the delta lineage. In addition, this PCLs triplet is found in four populations, two in Switzerland and two in Russia. This allelic consistency across lineages and populations suggests a conserved function that may contribute to broad host attachment. Along with being highly conserved, PCL53 seems to confer a nearly universal attachment (at the level of the host genotype panel used here, as well as a broader one, **Error! Reference source not found**.). While the benefits of PCL53 are clear, it is unclear why it is not found in many isolates. Having this gene might be costly, but what those costs are is unclear. It may reduce the infectivity of other PCL genes in the same isolate, reduce fitness for other traits (e.g. growth rate or competitive ability), or reduce the ability to infect other hosts. In addition, because Daphnia’s external abdomen consists of a thicker cuticle, penetrating into the host’s body cavity is probably harder for the parasite, making the rate of infection through this attachment site lower than at other sites (Fredericksen et al., 2021), indicating a reduced fitness. How the PCL53 presence/absence polymorphism contributes to the Red Queen dynamics of this system is unclear.

### Multiple proteins to attach on the same site?

In beta isolates, foregut attachment to the RU-C20-1 host appeared to be associated with PCL07, specifically with alleles A3, A5, and A7. However, alignment analysis did not reveal private SNPs specific to these alleles that could explain their role in attachment, a pattern also seen in attachment to the foregut of US-SP131-1 This raises several potential hypotheses: first, the association may be a false positive, so these alleles are not truly responsible for attachment. Alternatively, the same attachment site could be accessible to different proteins, thus explaining the difficulty of finding common SNPs.

### Lineage specific phenotypic effects

Our analysis assumed that single gene effects could explain *P. ramosa* phenotypes. However, an unexplained finding was the strong and specific attachment of gamma lineage isolates to the foregut of host clones US-SP131-1 and RU-C20-1; this attachment, unlike other observed patterns, does not appear to be driven by a single PCL allele or allele combination. To delineate this signal, we excluded the beta isolates from the GWAS but found that the signal in the PCL07 was lost. This suggests that the signal in PCL07 for foregut attachment on the host clone RU-C20-1 is driven by the beta isolates and leaves us without explanation for the attachment of the gamma isolate to the foregut of the host clones RU-C20-1 and US-SP131-1.

### Challenges in Detecting Presence-Absence Variations (PAVs)

A major challenge of our study was to overcome the limitations of SNP-based GWAS, which often fails to capture structural variants like presence-absence variations (PAVs) that may be essential to traits like host attachment. Because SNP-based GWAS traditionally relies on a single reference genome, it overlooks genetic elements that might be absent from or differ significantly in the reference. In contrast, our gene-PAV GWAS approach integrates reads that do not map to the reference genome, allowing us to detect genetic components that would otherwise be missed. We assembled non-host reads and filtered for contigs that aligned to at least one of three lineage-specific reference genomes, which we anticipated would encompass all PCLs to be detected in the present data set. While the short-read lengths generated from Illumina sequencing posed assembly limitations, we proceeded under the hypothesis that the presence of specific PCLs would be sufficient for host attachment. The assembled genomes had completeness scores ranging from 80% to 97.82%, allowing for a robust pangenome analysis across lineages. This approach allowed us to identify triplet PCL52-53-54 as a potential candidate for attachment to the external abdomen across most host clones.

## Conclusion

This study advances our understanding of the genetic mechanisms underlying host–parasite coevolution by examining the role of collagen-like protein (PCL) genes in the attachment phenotypes of *Pasteuria ramosa* to its host, *Daphnia magna*. By integrating genomic and phenotypic data, we uncovered significant diversity within the PCL gene family, highlighting its evolutionary importance. Specifically, our findings reveal lineage-specific patterns of PCL gene content and diversity, with distinct associations between certain PCL triplets and attachment phenotypes to host attachment sites. These results underscore the critical role of genetic variation in facilitating host–parasite interactions and support the Red Queen model as a driving force behind the observed diversity.

Notably, our study provides strong evidence for the matching-allele-model, particularly through our analyses of PCL22 and PCL07 where the specific effect of *P. ramosa* alleles is conditional to the host genotype. We observed strong genotype-phenotype segregation for PCL07 and PCL22, with specific alleles showing mutual exclusivity in driving attachment phenotypes to distinct host foregut genotypes. Replacing one allele with another can reverse the infection patterns, a hallmark of coevolutionary processes. However, such a reversal was not observed for a presence/absence polymorphism of PCL53 and a linked infection phenotype. Whether that is because we tested too few host genotypes, or because it does not exist is currently unclear.

Our results also demonstrate the utility of combining SNP-based and PAV GWAS to capture both subtle and structural genetic differences that influence infection outcomes. This dual approach allowed us to pinpoint several PCL triplets, such as PCL07, PCL22 and PCL53, as key players in host attachment, while also revealing lineage-specific differences and recombination events that further contribute to genetic diversity and adaptive potential.

By linking genetic diversity and genotype matching dynamics to host–parasite interactions, our study underscores the importance of genetic matching in facilitating coevolution. These findings provide critical insights into the molecular mechanisms of pathogen adaptation and highlight the evolutionary processes that sustain diversity in dynamic host–parasite systems. Beyond advancing theoretical frameworks like the Red Queen model, our results have broader implications for managing pathogen evolution and understanding genetic mechanisms in other coevolving systems.

## Material and Methods

### Collection and preparation of the *Pasteuria ramosa* diversity panel

Most isolates were isolated from pond sediment through different experimental infection as part of the previously reported population structure study (**Error! Reference source not found**.)(Andras et al., 2018). For this study, 34 further isolates were isolated from pond sediments following the same protocol.

### DNA extraction and whole genome sequencing

To obtain good quality DNA, we extracted the DNA while the parasite was still in its immature stage. The isolate was propagated in its host clone until two weeks after the infection, when we collected the infected host as described in (Dexter et al., 2023).

DNA was extracted from infected *Daphnia* using a modified protocol based on the Gentra PureGene extraction kit (Qiagen). In brief, whole *Daphnia* were homogenized with a pestle and incubated overnight at 55 °C in lysis buffer with Proteinase K. Following this, samples were treated with RNase A at 37 °C for 30 minutes, then protein was precipitated by centrifugation. DNA was precipitated by addition to chilled isopropanol and glycogen; after centrifugation, the DNA pellet was washed once with 70% ethanol. Finally, the DNA was resuspended in buffer solution and stored at ™20 °C.

Genomic libraries for short-read sequencing were prepared using a NEBNext Ultra II DNA Library Prep Kit for Illumina. The libraries were sequenced across multiple runs of a Novaseq 6000 to generate paired-end 100 bp reads aiming for at least 50× coverage across the parasite genomes. Genomic library preparation and Illumina sequencing were performed at the Basel Genomics Facility, Basel, Switzerland.

### SNP calling and GWAS

Read pairs were trimmed using Trimmomatic version 0.39 to remove low-quality reads and adaptor sequences (Bolger et al., 2014). The trimmed reads were then mapped against the C1 reference genome (GNKnumber) using BWA-MEM2 version 2.2.1 set to default parameters (Vasimuddin et al., 2019). Duplicate reads were removed using Picard version 2.26.10 (*Picard Toolkit*, 2019). We jointly called genomic variants (SNPs, indels, and mixed-type variants) across all samples using the GATK haplotype caller and following GATK Best Practices recommendations (Auwera & O’Connor, 2020). Double infections were identified and, when found, the sample was excluded. The VCF file was filtered to remove low-quality bases and poor alignment scores. Variants were filtered to exclude sites with excessive depth (>2× mean depth), and VCF files were filtered to remove multi-allelic variants (BCFtools v 1.15, (Danecek et al., 2021)).

Pixy (v1.2.7.beta1; (Korunes & Samuk, 2021)) was used to calculate nucleotide diversity (π) across the genome with 100 bp sliding windows, as well as for each PCL. Since not all PCLs are present in the C1 genome, we repeated SNP calling in the P54 reference genome to capture the full diversity of PCLs. For PCLs present in both genomes, we averaged a diversity score using the two measurements.

To prepare genomic data for GWAS, we applied the following filters in Plink2 (v2.00a2.3; (Chang et al., 2015)): --geno 0.90, --maf 0.10, --mac 10, and --mind 0.75. Linkage disequilibrium filtering was performed with --indep-pairwise 1kb 1 0.9. Isolates exhibiting signs of multiple infection (i.e high heterozygosity) were excluded, leaving 91 total isolates. GWAS analysis was done with Plink2, using an infectotype matrix with a 50% cut-off. We accounted for lineage effects in the dataset by adding the lineages and sub-lineages as covariates. A similar result is obtained by correcting with the 4 first principal components generated by Plink2 from the genotype data, representing 72% of variance.

For admixture analysis, we used the same SNP set as for GWAS, with cross-validation applied to select the optimal value of K, ranging from 1 to 10 (v1.3.0; (Alexander et al., 2009)). Values of K greater than 4 indicated within-lineage clustering rather than lineage-level signals and were thus excluded. Principal Components Analysis (PCA) was conducted on the same dataset using Plink.

To infer the phylogenetic tree, the remaining 6,149 genome-wide SNPs were used. For consistency among the different figures containing the phylogenetic tree, we used isolates that had a completeness over 80 %, were stick tested, and showed no signs of double infection. 91 isolates fit these requirements. The best-fit model was identified as SYM+ASC+G4 using IQ-TREE (IQ tree v2.0, (Kalyaanamoorthy et al., 2017; Minh et al., 2020)). The tree was then computed with RAxML-NG (v 1.1 (Kozlov et al., 2019)).

### De novo assembly, PCLs catalog and alleles

To maximize the recovery of PCLs, all genomes were assembled. Paired reads that did not map to *Daphnia* (gene bank ID, BWA-MEM2 version 2.2.1 (Vasimuddin et al., 2019)) were extracted and assembled using MEGAHIT (version 1.2.9, (D. Li et al., 2015)) with default parameters. These assemblies were then annotated with Prokka (v1.14.5; (Seemann, 2014)), and PCLs were detected using the HMM profile PFM01391 in HMMER (v3.3.2; (Eddy, 2011)). Identified PCLs were matched to previously cataloged PCLs using BLAST+ (v2.13.0; (Camacho et al., 2009)), with PCLs showing over 70% homology to known sequences being named accordingly. Distinct PCLs were aligned with known PCLs, and an ML tree was generated (data not shown) to cluster new PCLs within established groups (groups 1, 2, 3, and 4; (McElroy et al., 2011)), resulting in the naming of clusters such as PCL54-53-52, PCL55-56-57, and PCL60-59-58.

For each PCL we used Clustal Omega (v 1.2.4, (Sievers et al., 2011)) to conduct a multi pairwise alignment and generated the Pairwise Identity Matrice (PIM) using Clustaw2 (v 2.1, (Larkin et al., 2007)). All proteins with over 95% homology were clustered in one allele group. PCLs/ alleles with a size difference of 10% were classified as N terminal/C terminal fragments or just fragments.

### Presence absence variation analysis

To retain primarily *Pasteuria*-related contigs and genes, assembled contigs were aligned to three reference genomes representing the three major lineages (GNBKnumber, C1, P54, and P21) with minimap2 (v 2.20, (H. Li, 2018)). Only contigs mapping to at least one of the three reference genomes and the corresponding annotations were retained. Genomes with a completeness higher than 80% were retained. PPanGGolin (v 2.2.0, (Gautreau et al., 2021) was then run on the GFF files using default settings.

To link genes with traits, Scoary2 (v 0.0.15,(Roder et al., 2024) was applied to the presence-absence matrix generated by PPanGGolin and the infectotype matrix previously binarized with a cutoff at 50%.

### Phenotyping of the isolates

Attachment phenotypes were scored according to the methods of (Ameline et al., 2022; Duneau et al., 2011; Fredericksen et al., 2021). In brief, three-to-five-day-old Daphnia are put in the presence of 10,000 fluorescent spores in individual wells containing 150 µL Daphnia medium (=ADaM). After 1 h of incubation in the dark, each well is rinsed twice to remove excess spores, and individual Daphnia are scored for attachment at each host position--the foregut (F); the hindgut (subdivided into three sites, the rectum (R), distal hindgut (D), and anus (A)); the external abdomen (E); the fourth (L4) and fifth (L5) trunk limb, and all trunk limbs (LA). A minimum of six Daphnia per combination host genotype x isolate were tested (i.e. replicates). Most *P. ramosa* isolates were assessed for their attachment phenotypes by researcher AT (**Error! Reference source not found**.), the other data was taken from Fredericksen et al. (2021).

## Data Availability Statement

Raw data is deposited at the NCBI SRA database (XXXXX). Scripts are available at https://github.com/AThivolle/XXX/.

They will be publicly available upon acceptance.

## Author Contributions

All authors designed the study. AT performed the molecular work. AT analyzed the data. AT and DE wrote the manuscript. AT and DE reviewed the manuscript.

## Acknowledgements

We thank Jürgen Hottinger, Urs Stiefel and Michelle Krebs for help in the laboratory. We thank members of the Ebert group for providing feedback on the study and the manuscript.

## References

Agrawal, A., & Lively, C. M. (2002). Infection genetics: Gene-for-gene versus matching-alleles models and all points in between. Evolutionary Ecology Research, 4(1), 79–90.

Akther, S., Mongodin, E. F., Morgan, R. D., Di, L., Yang, X., Golovchenko, M., Rudenko, N., Margos, G., Hepner, S., Fingerle, V., Kawabata, H., Norte, A. C., de Carvalho, I. L., Núncio, M. S., Marques, A., Schutzer, S. E., Fraser, C. M., Luft, B. J., Casjens, S. R., & Qiu, W. (2024). Natural selection and recombination at host-interacting lipoprotein loci drive genome diversification of Lyme disease and related bacteria. mBio, 15(9), e01749–24. 10.1128/mbio.01749-24

Alexander, D. H., Novembre, J., & Lange, K. (2009). Fast model-based estimation of ancestry in unrelated individuals. Genome Research, 19(9), 1655–1664. 10.1101/gr.094052.109

Ameline, C., Voegtli, F., Jason, A., Dexter, E., Engelstädter, J., & Dieter, E. (2022). Genetic slippage after sex maintains diversity for parasite resistance in a natural host population. SCIENCE ADVANCES. 10.1126/sciadv.abn0051

Andras, J. P., Fields, P. D., Du Pasquier, L., Fredericksen, M., & Ebert, D. (2020). Genome-wide association analysis identifies a genetic basis of infectivity in a model bacterial pathogen. Molecular Biology and Evolution. 10.1093/molbev/msaa173

Andras, J. P., Fields, P. D., & Ebert, D. (2018). Spatial population genetic structure of a bacterial parasite in close coevolution with its host. Molecular Ecology, 27(6), 1371–1384. 10.1111/mec.14545

Arnold, B. J., Huang, I.-T., & Hanage, W. P. (2022). Horizontal gene transfer and adaptive evolution in bacteria. Nature Reviews Microbiology, 20(4), 206–218. 10.1038/s41579-021-00650-4

Auwera, G. V. der, & O’Connor, B. D. (2020). Genomics in the cloud: Using Docker, GATK, and WDL in Terra (First edition). O’Reilly.

Bachert, B. A., Choi, S. J., Snyder, A. K., Rio, R. V. M., Durney, B. C., Holland, L. A., Amemiya, K., Welkos, S. L., Bozue, J. A., Cote, C. K., Berisio, R., & Lukomski, S. (2015). A Unique Set of the Burkholderia Collagen-Like Proteins Provides Insight into Pathogenesis, Genome Evolution and Niche Adaptation, and Infection Detection. PLOS ONE, 10(9). 10.1371/journal.pone.0137578

Bolger, A. M., Lohse, M., & Usadel, B. (2014). Trimmomatic: A flexible trimmer for Illumina sequence data. Bioinformatics, 30(15), 2114–2120. 10.1093/bioinformatics/btu170

Camacho, C., Coulouris, G., Avagyan, V., Ma, N., Papadopoulos, J., Bealer, K., & Madden, T. L. (2009). BLAST+: Architecture and applications. BMC Bioinformatics, 10(1), 421. 10.1186/1471-2105-10-421

Carius, H., Little, T., & Ebert, D. (2001). Genetic variation in a host-parasite association: Potential for coevolution and frequency dependent selection. Evolution, 55(6), 1136–1145. http://www.jstor.org/stable/2680280 %5Cnpapers2://publication/uuid/50BE97C5-83D7-41FA-8BE5-AB8A03C7FF58

Chang, C. C., Chow, C. C., Tellier, L. C., Vattikuti, S., Purcell, S. M., & Lee, J. J. (2015). Second-generation PLINK: Rising to the challenge of larger and richer datasets. GigaScience, 4(1), 7. 10.1186/s13742-015-0047-8

Charlesworth, D. (2006). Balancing Selection and Its Effects on Sequences in Nearby Genome Regions. PLoS Genetics, 2(4), e64. 10.1371/journal.pgen.0020064

Chen, L., Mathema, B., Pitout, J. D. D., DeLeo, F. R., & Kreiswirth, B. N. (2014). Epidemic Klebsiella pneumoniae ST258 Is a Hybrid Strain. mBio, 5(3), 10.1128/mbio.01355-14. 10.1128/mbio.01355-14

Danecek, P., Bonfield, J. K., Liddle, J., Marshall, J., Ohan, V., Pollard, M. O., Whitwham, A., Keane, T., McCarthy, S. A., Davies, R. M., & Li, H. (2021). Twelve years of SAMtools and BCFtools. GigaScience, 10(2), giab008. 10.1093/gigascience/giab008

Decaestecker, E., Gaba, S., Raeymaekers, J. A. M., Stoks, R., Van Kerckhoven, L., Ebert, D., & De Meester, L. (2007). Host-parasite “Red Queen” dynamics archived in pond sediment. Nature, 450(7171), 870–873. 10.1038/nature06291

Dexter, E., Fields, P. D., & Ebert, D. (2023). Uncovering the Genomic Basis of Infection Through Co-genomic Sequencing of Hosts and Parasites. Molecular Biology and Evolution, Volume 40(Issue 7). 10.1093/molbev/msad145

Duneau, D., Luijckx, P., Ben-Ami, F., Laforsch, C., & Ebert, D. (2011). Resolving the infection process reveals striking differences in the contribution of environment, genetics and phylogeny to host-parasite interactions. BMC Biology, 9(February). 10.1186/1741-7007-9-11

Dutta, C., & Pan, A. (2002). Horizontal gene transfer and bacterial diversity. Journal of Biosciences, 27(1 SUPPL. 1), 27–33. 10.1007/BF02703681

Ebert, D. (1996). Development, life cycle, ultrastructure and phylogenetic position of Pasteuria ramosa Metchnikoff 1888: Rediscovery of an obligate endoparasite of Daphnia magna Straus. Philosophical Transactions of the Royal Society B: Biological Sciences, 351(1348), 1689–1701. 10.1098/rstb.1996.0151

Ebert, D., & Fields, P. D. (2020). Host–parasite co-evolution and its genomic signature. Nature Reviews Genetics, 21(12), 754–768. 10.1038/s41576-020-0269-1

Eddy, S. R. (2011). Accelerated Profile HMM Searches. PLoS Computational Biology, 7(10), e1002195. 10.1371/journal.pcbi.1002195

Frantz, C., Stewart, K. M., & Weaver, V. M. (2010). The extracellular matrix at a glance. Journal of Cell Science, 123(24), 4195–4200. 10.1242/jcs.023820

Fredericksen, M., Ameline, C., Krebs, M., Hüssy, B., Fields, P. D., Andras, J. P., & Ebert, D. (2021). Infection phenotypes of a coevolving parasite are highly diverse, structured, and specific. Evolution, 75(10), 2540–2554. 10.1111/evo.14323

Fredericksen, M., Fields, P. D., Du Pasquier, L., Ricci, V., & Ebert, D. (2023). QTL study reveals candidate genes underlying host resistance in a Red Queen model system. PLOS Genetics, 19(2), e1010570. 10.1371/journal.pgen.1010570

Gautreau, G., Bazin, A., Gachet, M., Planel, R., Burlot, L., Dubois, M., Perrin, A., Médigue, C., Calteau, A., Cruveiller, S., Matias, C., Ambroise, C., Rocha, E. P. C., & Vallenet, D. (2021). PPanGGOLiN: Depicting microbial diversity via a partitioned pangenome graph. PLOS Computational Biology, 17(12), e1009687. 10.1371/journal.pcbi.1009687

Hamilton, W. D. (1980). Sex versus Non-Sex versus Parasite. Wiley on Behalf of Nordic Society Oikos, 35(2), 282–290. http://www.jstor.org/stable/3544435

Huessy, B., Bumann, D., & Ebert, D. (2023). Ectopical expression of bacterial collagen-like protein supports its role as adhesin in host-parasite coevolution [Preprint]. Evolutionary Biology. 10.1101/2023.07.14.549037

Jackson, L. K., Potter, B., Schneider, S., Fitzgibbon, M., Blair, K., Farah, H., Krishna, U., Bedford, T., Peek, R. M., & Salama, N. R. (2020). Helicobacter pylori diversification during chronic infection within a single host generates sub-populations with distinct phenotypes. PLOS Pathogens, 16(12), e1008686. 10.1371/journal.ppat.1008686

Kalyaanamoorthy, S., Minh, B. Q., Wong, T. K. F., Von Haeseler, A., & Jermiin, L. S. (2017). ModelFinder: Fast model selection for accurate phylogenetic estimates. Nature Methods, 14(6), 587–589. 10.1038/nmeth.4285

Kirkman, L. A., & Deitsch, K. W. (2014). Recombination and Diversification of the Variant Antigen Encoding Genes in the Malaria Parasite Plasmodium falciparum. Microbiology Spectrum, 2(6), 10.1128/microbiolspec.mdna3-0022–2014. 10.1128/microbiolspec.mdna3-0022-2014

Korunes, K. L., & Samuk, K. (2021). PIXY: Unbiased estimation of nucleotide diversity and divergence in the presence of missing data. Molecular Ecology Resources, 21(4), 1359–1368. 10.1111/1755-0998.13326

Kozlov, A. M., Darriba, D., Flouri, T., Morel, B., & Stamatakis, A. (2019). RAxML-NG: a fast, scalable and user-friendly tool for maximum likelihood phylogenetic inference. Bioinformatics, 35(21), 4453–4455. 10.1093/bioinformatics/btz305

Larkin, M. A., Blackshields, G., Brown, N. P., Chenna, R., McGettigan, P. A., McWilliam, H., Valentin, F., Wallace, I. M., Wilm, A., Lopez, R., Thompson, J. D., Gibson, T. J., & Higgins, D. G. (2007). Clustal W and Clustal X version 2.0. Bioinformatics, 23(21), 2947–2948. 10.1093/bioinformatics/btm404

Li, D., Liu, C.-M., Luo, R., Sadakane, K., & Lam, T.-W. (2015). MEGAHIT: An ultra-fast single-node solution for large and complex metagenomics assembly via succinct de Bruijn graph. Bioinformatics, 31(10), 1674–1676. 10.1093/bioinformatics/btv033

Li, H. (2018). Minimap2: Pairwise alignment for nucleotide sequences. Bioinformatics, 34(18), 3094–3100. 10.1093/bioinformatics/bty191

Lively, C. M. (2010). A review of red queen models for the persistence of obligate sexual reproduction. Journal of Heredity, 101(SUPPL. 1), 13–20. 10.1093/jhered/esq010

Luijckx, P., Ben-Ami, F., Mouton, L., Du Pasquier, L., & Ebert, D. (2011). Cloning of the unculturable parasite Pasteuria ramosa and its Daphnia host reveals extreme genotype-genotype interactions. Ecology Letters, 14(2), 125–131. 10.1111/j.1461-0248.2010.01561.x

Luijckx, P., Fienberg, H., Duneau, D., & Ebert, D. (2013). A matching-allele model explains host resistance to parasites. Current Biology, 23(12), 1085–1088. 10.1016/j.cub.2013.04.064

Lukomski, S., Bachert, B. A., Squeglia, F., & Berisio, R. (2017). Collagen-like proteins of pathogenic streptococci. Molecular Microbiology, 103(6). 10.1111/mmi.13604

Lukomski, S., Nakashima, K., Abdi, I., Cipriano, V. J., Ireland, R. M., Reid, S. D., Adams, G. G., & Musser, J. M. (2000). Identification and Characterization of the scl Gene Encoding a Group A Streptococcus Extracellular Protein Virulence Factor with Similarity to Human Collagen. Infection and Immunity, 68(12), 6542–6553. 10.1128/IAI.68.12.6542-6553.2000

McElroy, K., Mouton, L., Du Pasquier, L., Qi, W., & Ebert, D. (2011). Characterisation of a large family of polymorphic collagen-like proteins in the endospore-forming bacterium Pasteuria ramosa. Research in Microbiology, 162(7), 701–714. 10.1016/j.resmic.2011.06.009

Minh, B. Q., Schmidt, H. A., Chernomor, O., Schrempf, D., Woodhams, M. D., Von Haeseler, A., & Lanfear, R. (2020). IQ-TREE 2: New Models and Efficient Methods for Phylogenetic Inference in the Genomic Era. Molecular Biology and Evolution, 37(5), 1530–1534. 10.1093/molbev/msaa015

Nielsen, R. (2001). Statistical tests of selective neutrality in the age of genomics. Heredity, 86(6), 641–647. 10.1046/j.1365-2540.2001.00895.x

Nuismer, S. L., Jenkins, C. E., & Dybdahl, M. F. (2017). Identifying coevolving loci using interspecific genetic correlations. Ecology and Evolution, 7(17), 6894–6903. 10.1002/ece3.3107

Oliver-Kozup, H. A., Elliott, M., Bachert, B. A., Martin, K. H., Reid, S. D., Schwegler-Berry, D. E., Green, B. J., & Lukomski, S. (2011). The streptococcal collagen-like protein-1 (Scl1) is a significant determinant for biofilm formation by group a Streptococcus. BMC Microbiology, 11(1), 262. 10.1186/1471-2180-11-262

Picard toolkit. (2019). [Computer software]. Broad Institute. https://broadinstitute.github.io/picard/

Pizarro-Guajardo, M., Olguín-Araneda, V., Barra-Carrasco, J., Brito-Silva, C., Sarker, M. R., & Paredes-Sabja, D. (2014). Characterization of the collagen-like exosporium protein, BclA1, of Clostridium difficile spores. Anaerobe, 25, 18–30. 10.1016/j.anaerobe.2013.11.003

Qiu, Y., Zhai, C., Chen, L., Liu, X., & Yeo, J. (2021). Current Insights on the Diverse Structures and Functions in Bacterial Collagen-like Proteins. ACS Biomaterials Science & Engineering. 10.1021/acsbiomaterials.1c00018

Reid, K. B. M. (1993). Structure/function relationships in the collectins (mammalian lectins containing collagen-like regions). Biochemical Society Transactions, 21(2), 464–468. 10.1042/bst0210464

Roder, T., Pimentel, G., Fuchsmann, P., Stern, M. T., Von Ah, U., Vergères, G., Peischl, S., Brynildsrud, O., Bruggmann, R., & Bär, C. (2024). Scoary2: Rapid association of phenotypic multi-omics data with microbial pan-genomes. Genome Biology, 25(1), 93. 10.1186/s13059-024-03233-7

Seemann, T. (2014). Prokka: Rapid prokaryotic genome annotation. Bioinformatics (Oxford, England), 30(14), 2068–2069. 10.1093/bioinformatics/btu153

Sievers, F., Wilm, A., Dineen, D., Gibson, T. J., Karplus, K., Li, W., Lopez, R., McWilliam, H., Remmert, M., Söding, J., Thompson, J. D., & Higgins, D. G. (2011). Fast, scalable generation of high-quality protein multiple sequence alignments using Clustal Omega. Molecular Systems Biology, 7(1), 539. 10.1038/msb.2011.75

Sironi, M., Cagliani, R., Forni, D., & Clerici, M. (2015). Evolutionary insights into host–pathogen interactions from mammalian sequence data. Nature Reviews Genetics, 16(4), 224–236. 10.1038/nrg3905

Suerbaum, S., & Ailloud, F. (2023). Genome and population dynamics during chronic infection with Helicobacter pylori. Current Opinion in Immunology, 82, 102304. 10.1016/j.coi.2023.102304

Sylvestre, P., Couture-Tosi, E., & Mock, M. (2002). A collagen-like surface glycoprotein is a structural component of the Bacillus anthracis exosporium. Molecular Microbiology, 45(1), 169–178. 10.1046/j.1365-2958.2000.03000.x

Thivolle, A., Paljakka, M., Ebert, D., & Fields, P. D. (2024). The genome of Pasteuria ramosa reveals a high turnover rate of collagen-like genes (p. 2024.02.09.579640). bioRxiv. 10.1101/2024.02.09.579640

Vandersmissen, L., De Buck, E., Saels, V., Coil, D. A., & Anné, J. (2010). A Legionella pneumophila collagen-like protein encoded by a gene with a variable number of tandem repeats is involved in the adherence and invasion of host cells. FEMS Microbiology Letters, 306(2), 168–176. 10.1111/j.1574-6968.2010.01951.x

Vasimuddin, Md., Misra, S., Li, H., & Aluru, S. (2019). Efficient Architecture-Aware Acceleration of BWA-MEM for Multicore Systems. 2019 IEEE International Parallel and Distributed Processing Symposium (IPDPS), 314–324. 10.1109/IPDPS.2019.00041

Woods, L. C., Gorrell, R. J., Taylor, F., Connallon, T., Kwok, T., & McDonald, M. J. (2020). Horizontal gene transfer potentiates adaptation by reducing selective constraints on the spread of genetic variation. Proceedings of the National Academy of Sciences, 117(43), 26868–26875. 10.1073/pnas.2005331117

Woolhouse, M. E. J., Webster, J. P., Domingo, E., Charlesworth, B., & Levin, B. R. (2002). Biological and biomedical implications of the co-evolution of pathogens and their hosts. Nature Genetics, 32(4), 569–577. 10.1038/ng1202-569

