## supplementals for "Matching-Allele-Models explain host specificity and high diversity of Collagen-Like Proteins in a virulent pathogen"

### Supplementary

#### Figures

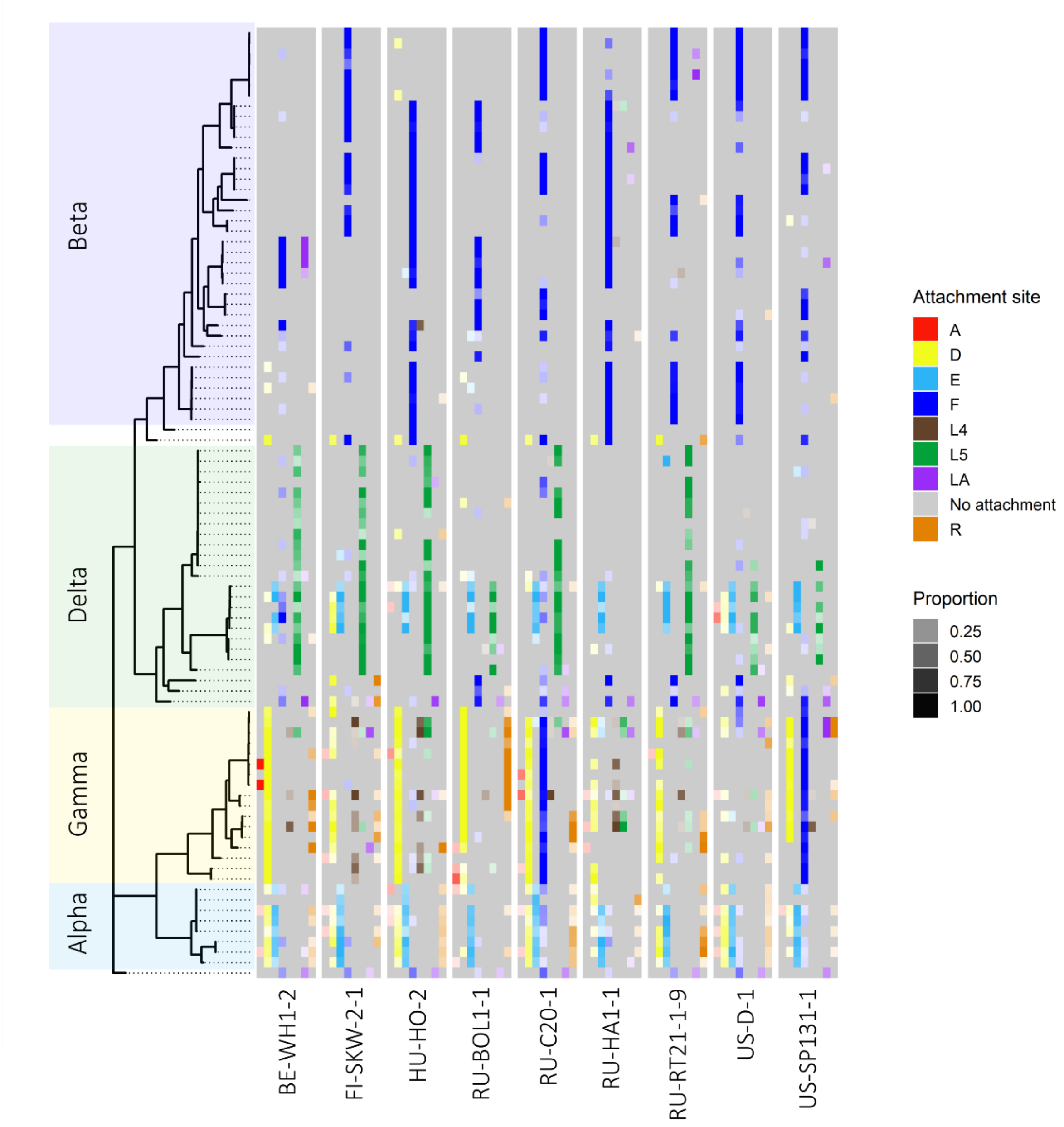


**Supplementary Figure 1 : Lineage-specific tropism for attachment sites.**
**Left Panel:** Phylogenetic tree as in Fig. 2.
**Right Panel:** Matrix representing the attachment of each isolate to each of the nine host panel clones. Attachment sites are color-coded, with color shades indicating the proportion of replicate attachments observed. Each isolate–host clone combination has been tested at least six times.

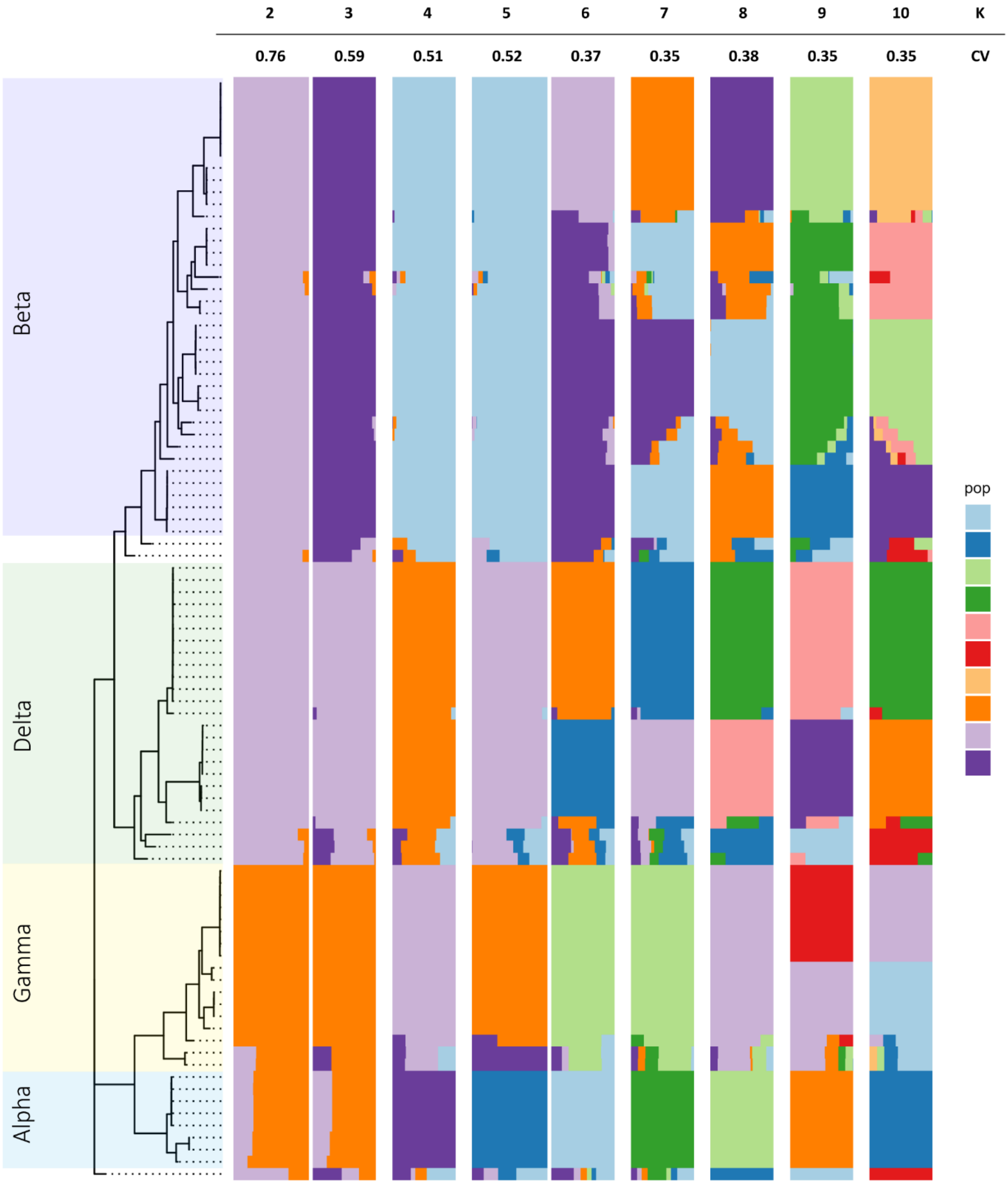


Supplementary Figure 2 : Admixture.

**Left Panel:** Phylogenetic tree as in Fig. 2.

**Right Panel:** Admixture results for K ranging from 2 to 10. A value of K=4 best captures the subdivision of *P. ramosa* in the four phylogenetic groups discussed here. A sub-population of the delta lineage shows a mix of ancestry, as well as a mix of PCLs and some phenotype deviation from the rest of the lineage.

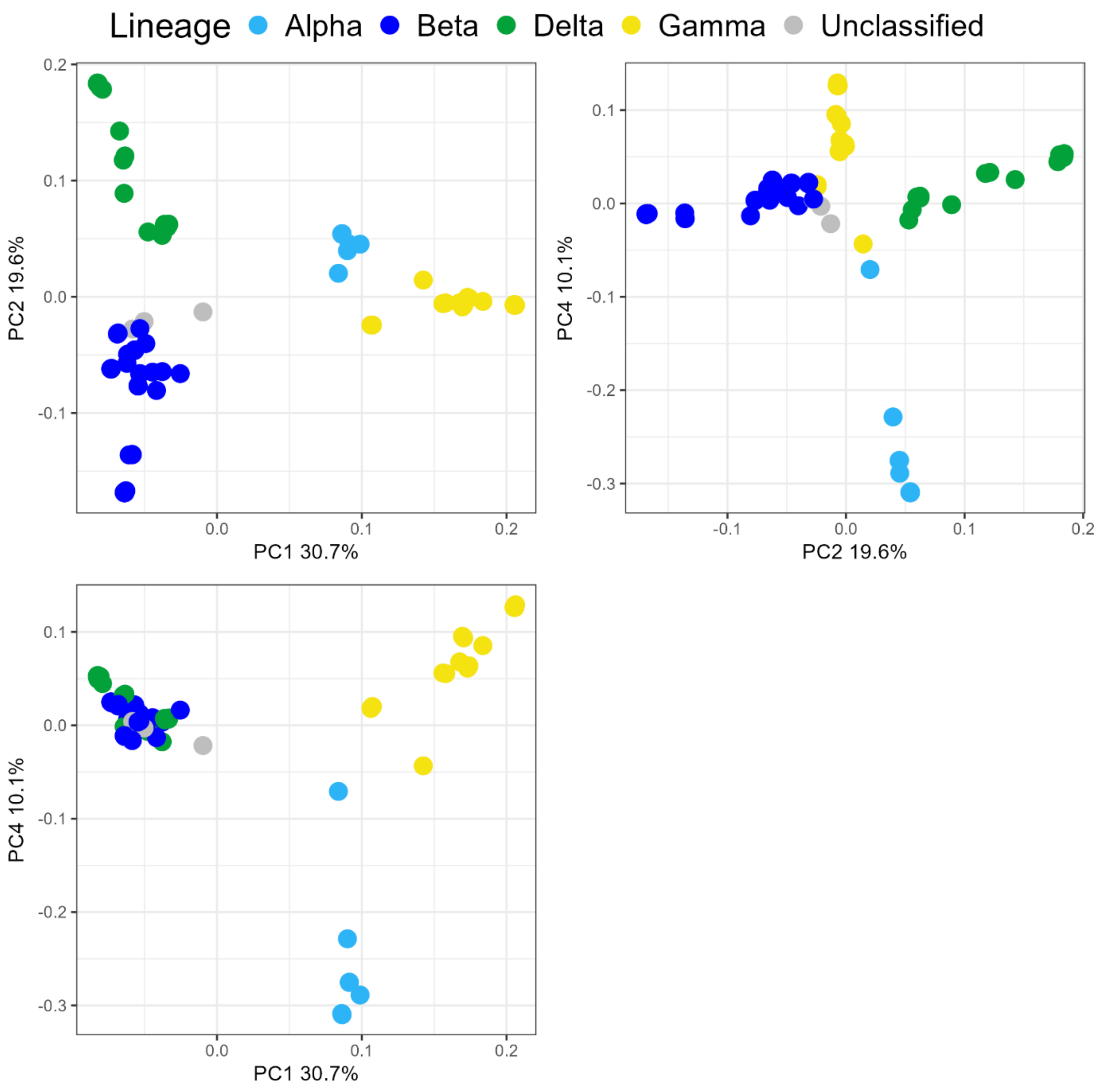


Supplementary Figure 3 : Principal component analysis (PCA) reveals strong data structure according to the *P. ramosa* lineage's structure.

PCA ordination of the four sub-lineages/lineages, PC1, PC2 and 4 allowed a neat separation of the sub-lineages. PC1 and PC2 together explain more than 50% of the variation.

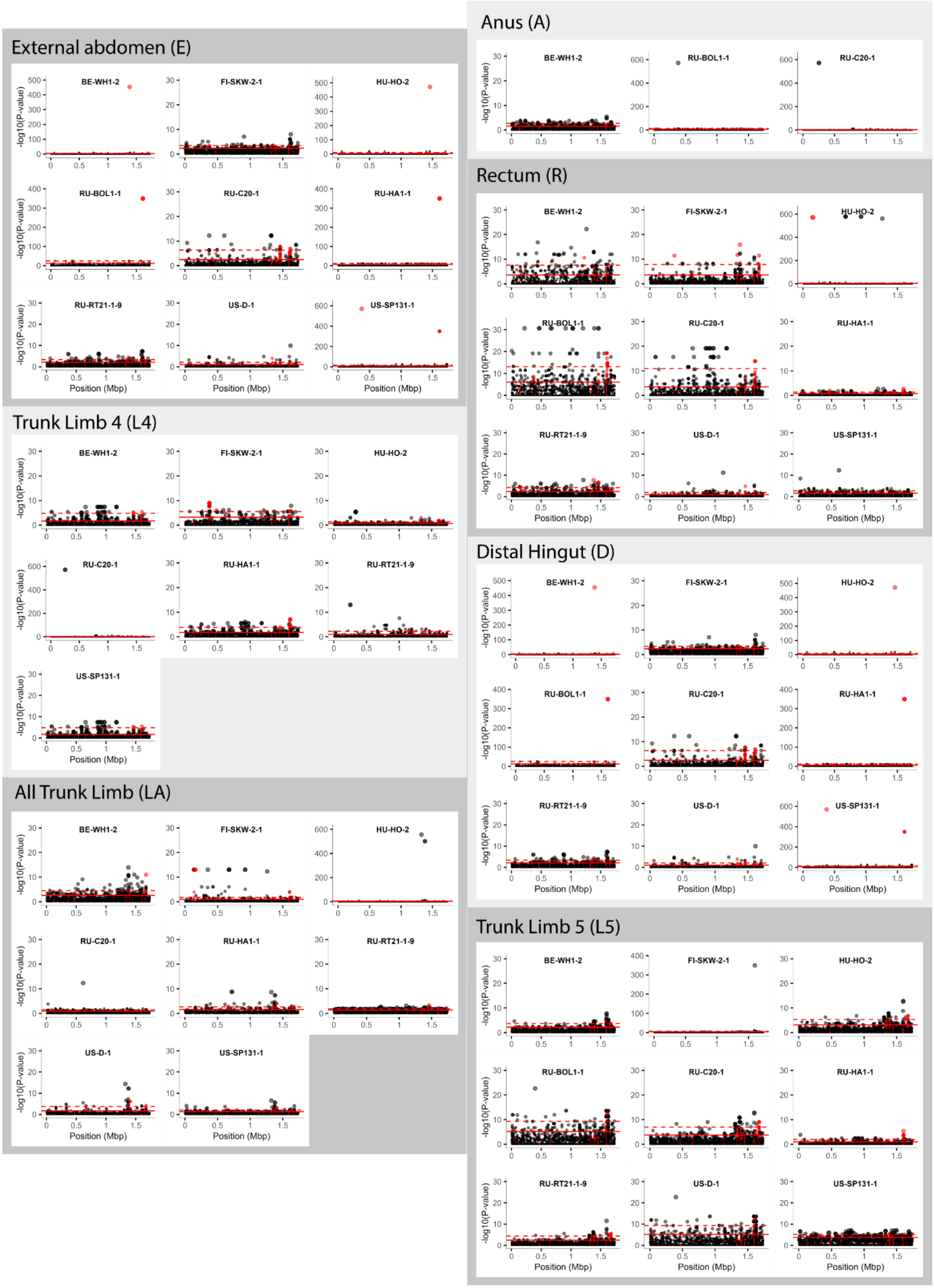


Supplementary Figure 4 : Manhattan plot of all attachment site, except the foregut (F) site, which is shown in Fig. 4.

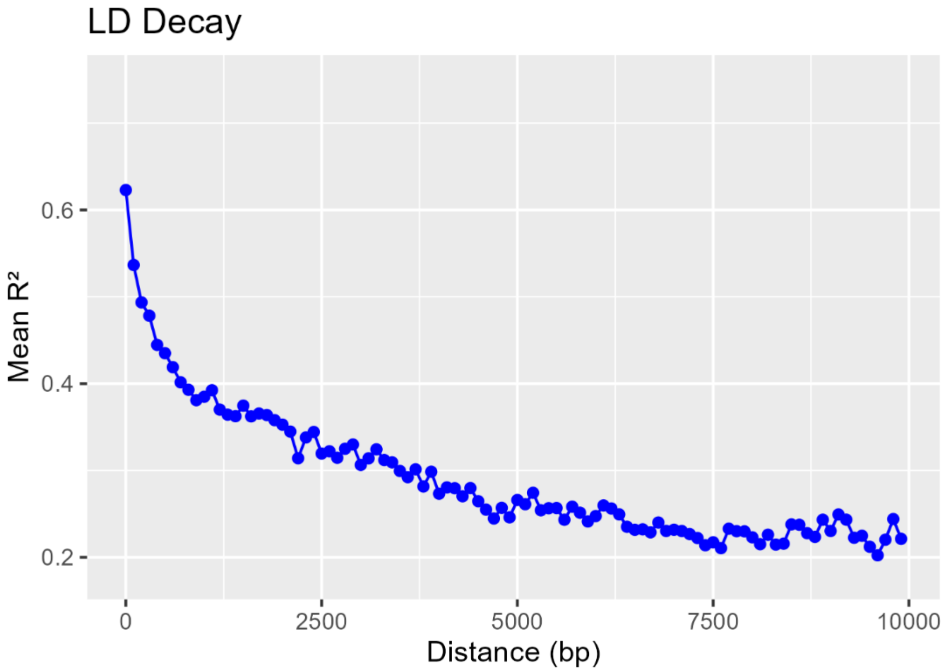


Supplementary Figure 5 : LD decay

Plot of linkage disequilibrium decay with distance across the *P. ramosa* genome for a distance of 10 kb. Pairwise LD was calculated for all variants across the genome and then mean values were calculated for 100 bp bins. Around a distance of 7.5 kb, the decay reaches the level of about 0.2, which is expected, given that the lineage structure of the samples implies some degree of LD across the entire genome.

### Tables

| Isolate name | Sediment infection (Study/N) | Locality | Latitude | Longitude | Sequencing |
|---|---|---|---|---|---|
| *B8_1* | This study | GB-S8-C | 51.621243 | -1.376437 | This study |
| *C1* | N | RU-RM1 | 55.763 | 37.582 | Andras et al 2020 |
| *C19* | N | DE-G1 | 54.282 | 10.966 | Andras et al 2020 |
| *C5_1* | This study | CH-Z-A | 47.51696 | 8.831272 | This study |
| *E2_1* | This study | BE-KN-B | 51.356787 | 3.342799 | This study |
| *G2_1* | This study | BE-KN-B | 51.356787 | 3.342799 | This study |
| *G2_2* | This study | BE-KN-B | 51.356787 | 3.342799 | This study |
| *G7_1* | This study | FI-SEG-2-A | 59.7642 | 23.3746 | This study |
| *H5_1* | This study | CH-Z-A | 47.51696 | 8.831272 | This study |
| *H6_1* | This study | CZ-KO-A1 | 50.12536111 | 14.86866667 | This study |
| *H8_1* | This study | GB-S8-C | 51.621243 | -1.376437 | This study |
| *H8_2* | This study | GB-S8-C | 51.621243 | -1.376437 | This study |
| *J2_1* | This study | BE-KN-B | 51.356787 | 3.342799 | This study |
| *J7_1* | This study | FI-SEG-2-A | 59.7642 | 23.3746 | This study |
| *J7_2* | This study | FI-SEG-2-A | 59.7642 | 23.3746 | This study |
| *J8_1* | This study | GB-S8-C | 51.621243 | -1.376437 | This study |
| *J8_2* | This study | GB-S8-C | 51.621243 | -1.376437 | This study |
| *K6_1* | This study | CZ-KO-A1 | 50.12536111 | 14.86866667 | This study |
| *K6_2* | This study | CZ-KO-A1 | 50.12536111 | 14.86866667 | This study |
| *K7_1* | This study | FI-SEG-2-A | 59.7642 | 23.3746 | This study |
| *K7_2* | This study | FI-SEG-2-A | 59.7642 | 23.3746 | This study |
| *K8_2* | This study | GB-S8-C | 51.621243 | -1.376437 | This study |
| *L6_1* | This study | CZ-KO-A1 | 50.12536111 | 14.86866667 | This study |
| *L6_2* | This study | CZ-KO-A1 | 50.12536111 | 14.86866667 | This study |
| *M7_2* | This study | FI-SEG-2-A | 59.7642 | 23.3746 | This study |
| *N5_1* | This study | CH-Z-A | 47.51696 | 8.831272 | This study |
| *N5_2* | This study | CH-Z-A | 47.51696 | 8.831272 | This study |
| *N8_1* | This study | GB-S8-C | 51.621243 | -1.376437 | This study |
| *O2_1* | This study | BE-KN-B | 51.356787 | 3.342799 | This study |
| *P1006* | Andras et al 2017 | RU-BN1 | 50.155519 | 43.3894 | This study |
| *P1012* | Andras et al 2017 | PL-1 | 52.21092 | 20.99654 | This study |
| *P1013* | Andras et al 2017 | PL-1 | 52.21092 | 20.99654 | Andras et al 2020 |
| *P1014* | Andras et al 2017 | PL-1 | 52.21092 | 20.99654 | This study |
| *P1015* | Andras et al 2017 | PL-1 | 52.21092 | 20.99654 | This study |
| *P1017* | Andras et al 2017 | RU-SYR1 | 53.212919 | 48.488211 | This study |
| *P1020* | Andras et al 2017 | CZ-1 | 50.12536111 | 14.86866667 | This study |
| *P1022* | Andras et al 2017 | CZ-1 | 50.12536111 | 14.86866667 | This study |
| *P1023* | Andras et al 2017 | EA1 | 51.877887 | -1.115456 | This study |
| *P1027* | Andras et al 2017 | GB-S17 | 51.620456 | -1.386222 | This study |
| *P1028* | Andras et al 2017 | GB-S17 | 51.620456 | -1.386222 | This study |
| *P1029* | Andras et al 2017 | BE-KN | 51.356787 | 3.342799 | Andras et al 2020 |
| *P1036* | Andras et al 2017 | FI-SEG1 | 59.7644 | 23.3742 | Andras et al 2020 |

| | | | | | |
|---|---|---|---|---|---|
| *P1042* | Andras et al 2017 | DE-L1 | 54.22214 | 10.4288 | Andras et al 2020 |
| *P1044* | Andras et al 2017 | DE-KN1 | 54.176906 | 10.806683 | Andras et al 2020 |
| *P15* | N | BE-OM2 | 50.863 | 4.721 | Andras et al 2020 |
| *P2* | N | GB-EK1 | 55.702406 | -2.340828 | This study |
| *P20* | N | CH-H | 47.558 | 8.863 | Andras et al 2020 |
| *P2009* | Andras et al 2017 | RU-SYR1 | 53.212919 | 48.488211 | Andras et al 2020 |
| *P2011* | Andras et al 2017 | RU-SYR1 | 53.212919 | 48.488211 | This study |
| *P2012* | Andras et al 2017 | RU-SYR1 | 53.212919 | 48.488211 | This study |
| *P2022* | Andras et al 2017 | FI-SEG1 | 59.7644 | 23.3742 | Andras et al 2020 |
| *P2025* | Andras et al 2017 | FI-SEG2 | 59.7642 | 23.3746 | This study |
| *P2031* | Andras et al 2017 | DE-KN1 | 54.176906 | 10.806683 | Andras et al 2020 |
| *P2037* | Andras et al 2017 | GB-EK1 | 55.702406 | -2.340828 | This study |
| *P2040* | Andras et al 2017 | GB-S8 | 51.621243 | -1.376437 | Andras et al 2020 |
| *P22* | N | RU-RM1 | 55.763472 | 37.581583 | This study |
| *P23* | N | RU-RM1 | 55.763472 | 37.581583 | This study |
| *P25* | N | FI-VIW5 | 59.8302 | 23.201017 | This study |
| *P28* | N | GB-EK1 | 55.702406 | -2.340828 | This study |
| *P3* | N | FI-VIW-2 | 59.83035833 | 23.20110278 | This study |
| *P30* | N | DE-G1 | 54.281929 | 10.966506 | This study |
| *P3005* | Andras et al 2017 | RU-SYR1 | 53.212919 | 48.488211 | Andras et al 2020 |
| *P3009* | Andras et al 2017 | CZ-1 | 50.12536111 | 14.86866667 | This study |
| *P3010* | Andras et al 2017 | CZ-1 | 50.12536111 | 14.86866667 | Andras et al 2020 |
| *P3017* | Andras et al 2017 | DE-N1 | 54.30854 | 10.6249 | This study |
| *P3019* | Andras et al 2017 | DE-KN1 | 54.176906 | 10.806683 | Andras et al 2020 |
| *P3020* | Andras et al 2017 | BE-WH2 | 51.334779 | 3.348134 | Andras et al 2020 |
| *P3022* | Andras et al 2017 | RU-RM1 | 55.763472 | 37.581583 | This study |
| *P3025* | Andras et al 2017 | RU-RT1 | 45.221667 | 36.808333 | This study |
| *P3034* | Andras et al 2017 | CH-H | 47.557769 | 8.862608 | Andras et al 2020 |
| *P32* | N | DE-G1 | 54.281929 | 10.966506 | This study |
| *P33* | N | DE-G1 | 54.281929 | 10.966506 | This study |
| *P34* | N | DE-G1 | 54.281929 | 10.966506 | This study |
| *P35* | N | DE-G1 | 54.281929 | 10.966506 | This study |
| *P36* | N | DE-G1 | 54.281929 | 10.966506 | This study |
| *P37* | N | IL-NS | 31.724185 | 34.626269 | This study |
| *P38* | N | CH-H | 47.557769 | 8.862608 | This study |
| *P39* | N | CH-H | 47.557769 | 8.862608 | This study |
| *P40* | N | CH-H | 47.557769 | 8.862608 | This study |
| *P4002_C* | Andras et al 2017 | CH-H | 47.557769 | 8.862608 | This study |

| | | | | | |
|---|---|---|---|---|---|
| ***P4005_C*** | Andras et al 2017 | RU-BN1 | 50.155519 | 43.3894 | This study |
| ***P4007*** | Andras et al 2017 | RU-BN1 | 50.155519 | 43.3894 | This study |
| ***P4008*** | Andras et al 2017 | EA24 | | | This study |
| ***P4014*** | Andras et al 2017 | CZ-1 | 50.12536111 | 14.86866667 | This study |
| ***P4017*** | Andras et al 2017 | CZ-1 | 50.12536111 | 14.86866667 | This study |
| ***P4021*** | Andras et al 2017 | BE-KN2 | 51.355731 | 3.334453 | This study |
| ***P4025*** | Andras et al 2017 | FI-SEG2 | 59.7642 | 23.3746 | This study |
| ***P4030*** | Andras et al 2017 | DE-N1 | 54.30854 | 10.6249 | This study |
| ***P4032*** | Andras et al 2017 | GB-EK1 | 55.702406 | -2.340828 | This study |
| ***P4035*** | Andras et al 2017 | DE-R1 | 54.2076 | 10.4236 | Andras et al 2020 |
| ***P4048*** | Andras et al 2017 | RU-RM1 | 55.763472 | 37.581583 | Andras et al 2020 |
| ***P4049*** | Andras et al 2017 | RU-RM1 | 55.763472 | 37.581583 | This study |
| ***P4054*** | Andras et al 2017 | RU-RT1 | 45.221667 | 36.808333 | Andras et al 2020 |
| ***P4056*** | Andras et al 2017 | FI-VIW5 | 59.8302 | 23.201017 | This study |
| ***P4057*** | Andras et al 2017 | FI-VIW5 | 59.8302 | 23.201017 | Andras et al 2020 |
| ***P4058*** | Andras et al 2017 | FI-VIW5 | 59.8302 | 23.201017 | This study |
| ***P4059*** | Andras et al 2017 | GB-S8 | 51.621243 | -1.376437 | Andras et al 2020 |
| ***P4060*** | Andras et al 2017 | GB-S8 | 51.621243 | -1.376437 | This study |
| ***P4064*** | Andras et al 2017 | CH-H | 47.557769 | 8.862608 | Andras et al 2020 |
| ***P45*** | N | CH-H | 47.557769 | 8.862608 | This study |
| ***P48*** | N | CH-H | 47.557769 | 8.862608 | This study |
| ***P49*** | N | CH-H | 47.557769 | 8.862608 | This study |
| ***P52*** | N | CH-H | 47.557769 | 8.862608 | This study |
| ***P53*** | N | CH-H | 47.557769 | 8.862608 | This study |
| ***P54*** | N | CH-H | 47.557769 | 8.862608 | This study |
| ***P55*** | N | CH-H | 47.557769 | 8.862608 | This study |
| ***P57*** | N | CH-H | 47.557769 | 8.862608 | This study |
| ***P58*** | N | CH-H | 47.557769 | 8.862608 | This study |
| ***P6_1*** | This study | CZ-KO-A1 | 50.12536111 | 14.86866667 | This study |
| ***P6_2*** | This study | CZ-KO-A1 | 50.12536111 | 14.86866667 | This study |
| ***R7_1*** | This study | FI-SEG-2-A | 59.7642 | 23.3746 | This study |
| ***R7_2*** | This study | FI-SEG-2-A | 59.7642 | 23.3746 | This study |
| ***S7_1*** | This study | FI-SEG-2-A | 59.7642 | 23.3746 | This study |
| ***S7_2*** | This study | FI-SEG-2-A | 59.7642 | 23.3746 | This study |
| ***S8_1*** | This study | GB-S8-C | 51.621243 | -1.376437 | This study |

| Isolate name | Stick-test | C1 coverage | Assembled | Completness compare to C1 | Multiple infections (Y/N) |
|---|---|---|---|---|---|
| *B8_1* | This study | 162.6 | y | 95.4 | N |
| *C1* | Fredericksen et al 2021 | 12.1 | y | 60.0 | NA |
| *C19* | Fredericksen et al 2021 | 108.1 | y | 85.5 | N |
| *C5_1* | This study | 444.8 | y | 97.4 | N |
| *E2_1* | This study | 218.5 | y | 95.3 | N |
| *G2_1* | This study | 101.9 | y | 95.9 | N |
| *G2_2* | This study | 173.3 | y | 97.5 | N |
| *G7_1* | This study | 286.0 | y | 97.5 | N |
| *H5_1* | This study | 475.5 | y | 97.5 | N |
| *H6_1* | This study | 230.8 | y | 96.1 | N |
| *H8_1* | This study | 268.3 | y | 95.1 | N |
| *H8_2* | This study | 265.5 | y | 97.1 | N |
| *J2_1* | This study | 101.8 | y | 94.9 | N |
| *J7_1* | This study | 17.7 | N | NA | NA |
| *J7_2* | N | 227.7 | y | 97.7 | N |
| *J8_1* | This study | 105.4 | y | 95.1 | N |
| *J8_2* | This study | 179.5 | y | 97.1 | N |
| *K6_1* | This study | 142.8 | y | 96.1 | N |
| *K6_2* | This study | 135.6 | y | 96.1 | N |
| *K7_1* | This study | 429.3 | y | 97.7 | N |
| *K7_2* | This study | 210.1 | y | 97.5 | N |
| *K8_2* | This study | 317.4 | y | 97.1 | N |
| *L6_1* | This study | 138.2 | y | 96.0 | N |
| *L6_2* | This study | 211.1 | y | 96.0 | N |
| *M7_2* | N | 166.9 | y | 97.6 | N |
| *N5_1* | This study | 0.0 | N | NA | NA |
| *N5_2* | This study | 162.6 | y | 95.7 | N |
| *N8_1* | This study | 170.1 | y | 95.3 | N |
| *O2_1* | N | 179.7 | y | 95.4 | N |
| *P1006* | This study | 253.4 | y | 97.4 | Y |
| *P1012* | This study | 351.3 | y | 97.5 | N |
| *P1013* | Fredericksen et al 2021 | 107.2 | y | 88.8 | N |
| *P1014* | This study | 208.9 | y | 97.4 | Y |
| *P1015* | This study | 187.1 | y | 97.3 | N |
| *P1017* | This study | 175.6 | y | 95.9 | N |
| *P1020* | This study | 204.5 | y | 96.1 | N |
| *P1022* | This study | 273.9 | y | 96.0 | N |
| *P1023* | N | 397.0 | y | 97.4 | N |
| *P1027* | This study | 336.9 | y | 95.2 | N |
| *P1028* | This study | 548.2 | y | 95.2 | N |
| *P1029* | Fredericksen et al 2021 | 135.1 | y | 87.2 | N |
| *P1036* | Fredericksen et al 2021 | 85.9 | y | 84.8 | N |
| *P1042* | Fredericksen et al 2021 | 101.7 | y | 87.5 | N |
| *P1044* | Fredericksen et al 2021 | 44.8 | y | 73.4 | NA |
| *P15* | Fredericksen et al 2021 | 117.0 | y | 88.2 | N |
| *P2* | Fredericksen et al 2021 | 654.3 | y | 95.1 | N |
| *P20* | Fredericksen et al 2021 | 82.1 | y | 84.1 | N |

| | | | | | |
|---|---|---|---|---|---|
| *P2009* | Fredericksen et al 2021 | 42.2 | y | 75.3 | NA |
| *P2011* | This study | 197.2 | y | 95.8 | N |
| *P2012* | This study | 106.1 | y | 95.8 | N |
| *P2022* | Fredericksen et al 2021 | 96.4 | y | 85.3 | N |
| *P2025* | This study | 323.4 | y | 95.8 | N |
| *P2031* | Fredericksen et al 2021 | 66.1 | y | 76.9 | N |
| *P2037* | This study | 951.9 | y | 96.4 | N |
| *P2040* | Fredericksen et al 2021 | 33.0 | y | 68.6 | NA |
| *P22* | Fredericksen et al 2021 | 268.2 | y | 95.7 | N |
| *P23* | Fredericksen et al 2021 | 803.9 | y | 96.0 | N |
| *P25* | Fredericksen et al 2021 | 709.2 | y | 97.7 | N |
| *P28* | Fredericksen et al 2021 | 1038.6 | y | 95.2 | N |
| *P3* | Fredericksen et al 2021 | 313.8 | y | 97.7 | N |
| *P30* | Fredericksen et al 2021 | 658.0 | y | 97.7 | N |
| *P3005* | Fredericksen et al 2021 | 116.2 | y | 86.7 | N |
| *P3009* | This study | 228.6 | y | 96.3 | N |
| *P3010* | N | 134.3 | y | 86.9 | N |
| *P3017* | This study | 246.4 | y | 95.6 | N |
| *P3019* | Fredericksen et al 2021 | 96.6 | y | 82.9 | N |
| *P3020* | Fredericksen et al 2021 | 53.0 | y | 75.0 | N |
| *P3022* | This study | 794.7 | y | 95.7 | N |
| *P3025* | This study | 623.7 | y | 95.6 | N |
| *P3034* | N | 129.0 | y | 86.9 | N |
| *P32* | Fredericksen et al 2021 | 468.2 | y | 97.3 | N |
| *P33* | Fredericksen et al 2021 | 660.2 | y | 97.6 | N |
| *P34* | Fredericksen et al 2021 | 477.1 | y | 97.5 | N |
| *P35* | Fredericksen et al 2021 | 868.9 | y | 97.7 | N |
| *P36* | Fredericksen et al 2021 | 513.1 | y | 97.6 | N |
| *P37* | Fredericksen et al 2021 | 417.2 | y | 97.5 | N |
| *P38* | Fredericksen et al 2021 | 298.9 | y | 95.5 | N |
| *P39* | Fredericksen et al 2021 | 258.9 | y | 95.6 | N |
| *P40* | Fredericksen et al 2021 | 454.7 | y | 95.7 | N |
| *P4002_C* | This study | 698.5 | y | 97.4 | N |
| *P4005_C* | This study | 493.6 | y | 97.5 | Y |
| *P4007* | This study | 1649.8 | y | 95.8 | N |
| *P4008* | This study | 1114.4 | y | 97.5 | N |
| *P4014* | This study | 747.7 | y | 96.1 | N |
| *P4017* | This study | 974.9 | y | 96.0 | N |
| *P4021* | This study | 579.5 | y | 96.1 | N |
| *P4025* | This study | 1679.7 | y | 97.6 | N |
| *P4030* | This study | 640.0 | y | 96.2 | N |
| *P4032* | This study | 914.3 | y | 95.2 | N |
| *P4035* | Fredericksen et al 2021 | 99.3 | y | 83.4 | N |
| *P4048* | Fredericksen et al 2021 | 85.2 | y | 81.3 | N |
| *P4049* | This study | 440.5 | y | 97.4 | N |
| *P4054* | Fredericksen et al 2021 | 80.8 | y | 77.9 | N |
| *P4056* | This study | 517.0 | y | 97.6 | N |
| *P4057* | Fredericksen et al 2021 | 133.1 | y | 86.2 | N |
| *P4058* | This study | 368.3 | y | 97.8 | Y |
| *P4059* | Fredericksen et al 2021 | 94.2 | y | 84.2 | N |

| | | | | | |
|---|---|---|---|---|---|
| *P4060* | This study | 1094.7 | y | 95.2 | N |
| *P4064* | Fredericksen et al 2021 | 160.0 | y | 89.6 | N |
| *P45* | Fredericksen et al 2021 | 52.7 | y | 95.7 | N |
| *P48* | Fredericksen et al 2021 | 266.2 | y | 97.4 | Y |
| *P49* | Fredericksen et al 2021 | 937.6 | y | 95.7 | N |
| *P52* | Fredericksen et al 2021 | 1255.8 | y | 95.5 | N |
| *P53* | Fredericksen et al 2021 | 623.5 | y | 97.6 | Y |
| *P54* | Fredericksen et al 2021 | 1219.6 | y | 95.7 | Y |
| *P55* | Fredericksen et al 2021 | 791.2 | y | 95.7 | N |
| *P57* | Fredericksen et al 2021 | 775.4 | y | 95.6 | Y |
| *P58* | Fredericksen et al 2021 | 23.5 | N | NA | NA |
| *P6_1* | This study | 294.7 | y | 96.2 | N |
| *P6_2* | This study | 386.5 | y | 96.1 | N |
| *R7_1* | This study | 485.2 | y | 97.5 | N |
| *R7_2* | This study | 304.4 | y | 97.7 | N |
| *S7_1* | This study | 360.1 | y | 97.8 | N |
| *S7_2* | This study | 588.8 | y | 97.7 | N |
| *S8_1* | This study | 347.8 | y | 97.1 | N |

| Isolate name | Excluded from GWAS (Y/N) | Lineage | Phylogenetic tree (Y/N) |
|---|---|---|---|
| *B8_1* | N | Gamma | Y |
| *C1* | Y | Beta | N |
| *C19* | N | Beta | Y |
| *C5_1* | N | Beta | Y |
| *E2_1* | N | Gamma | Y |
| *G2_1* | N | Delta | Y |
| *G2_2* | N | Beta | Y |
| *G7_1* | N | Beta | Y |
| *H5_1* | N | Beta | Y |
| *H6_1* | N | Delta | Y |
| *H8_1* | N | Gamma | Y |
| *H8_2* | N | Beta | Y |
| *J2_1* | N | Gamma | Y |
| *J7_1* | Y | NA | N |
| *J7_2* | Y | Beta | N |
| *J8_1* | N | Gamma | Y |
| *J8_2* | N | Beta | Y |
| *K6_1* | N | Delta | Y |
| *K6_2* | N | Delta | Y |
| *K7_1* | N | Beta | Y |
| *K7_2* | N | Beta | Y |
| *K8_2* | N | Beta | Y |
| *L6_1* | N | Delta | Y |
| *L6_2* | N | Delta | Y |
| *M7_2* | Y | Beta | N |
| *N5_1* | Y | NA | N |
| *N5_2* | N | Alpha | Y |
| *N8_1* | N | Gamma | Y |

| | | | |
|---|---|---|---|
| ***O2_1*** | Y | Gamma | N |
| ***P1006*** | Y | NA | N |
| ***P1012*** | N | Beta | Y |
| ***P1013*** | N | Beta | Y |
| ***P1014*** | Y | NA | N |
| ***P1015*** | N | Beta | Y |
| ***P1017*** | N | Delta | Y |
| ***P1020*** | N | Delta | Y |
| ***P1022*** | N | Delta | Y |
| ***P1023*** | Y | Beta | N |
| ***P1027*** | N | Gamma | Y |
| ***P1028*** | N | Gamma | Y |
| ***P1029*** | N | Gamma | Y |
| ***P1036*** | N | Gamma | Y |
| ***P1042*** | N | Beta | Y |
| ***P1044*** | Y | Gamma | N |
| ***P15*** | N | Gamma | Y |
| ***P2*** | N | Gamma | Y |
| ***P20*** | N | Beta | Y |
| ***P2009*** | Y | Delta | N |
| ***P2011*** | N | Delta | Y |
| ***P2012*** | N | Delta | Y |
| ***P2022*** | N | Beta | Y |
| ***P2025*** | N | Gamma | Y |
| ***P2031*** | Y | Gamma | N |
| ***P2037*** | N | Beta | Y |
| ***P2040*** | Y | Gamma | N |
| ***P22*** | N | Delta | Y |
| ***P23*** | N | Delta | Y |
| ***P25*** | N | Beta | Y |
| ***P28*** | N | Gamma | Y |
| ***P3*** | N | Beta | Y |
| ***P30*** | N | Beta | Y |
| ***P3005*** | N | Delta | Y |
| ***P3009*** | N | Delta | Y |
| ***P3010*** | Y | Delta | N |
| ***P3017*** | N | Delta | Y |
| ***P3019*** | N | Delta | Y |
| ***P3020*** | N | Delta | N |
| ***P3022*** | N | Delta | Y |
| ***P3025*** | N | Delta | Y |
| ***P3034*** | Y | Alpha | N |
| ***P32*** | N | Beta | Y |
| ***P33*** | N | Beta | Y |
| ***P34*** | N | Beta | Y |
| ***P35*** | N | Beta | Y |
| ***P36*** | N | Beta | Y |
| ***P37*** | N | Beta | Y |
| ***P38*** | N | Alpha | Y |
| ***P39*** | N | Alpha | Y |

| | | | |
|---|---|---|---|
| *P40* | N | Alpha | Y |
| *P4002_C* | N | Beta | Y |
| *P4005_C* | Y | NA | N |
| *P4007* | N | Delta | Y |
| *P4008* | N | Beta | Y |
| *P4014* | N | Delta | Y |
| *P4017* | N | Delta | Y |
| *P4021* | N | Delta | Y |
| *P4025* | N | Beta | Y |
| *P4030* | N | Beta | Y |
| *P4032* | N | Gamma | Y |
| *P4035* | N | Beta | Y |
| *P4048* | N | Delta | Y |
| *P4049* | N | Beta | Y |
| *P4054* | N | Alpha | N |
| *P4056* | N | Beta | Y |
| *P4057* | N | Beta | Y |
| *P4058* | Y | NA | N |
| *P4059* | N | Gamma | Y |
| *P4060* | N | Gamma | Y |
| *P4064* | N | Beta | Y |
| *P45* | N | Alpha | Y |
| *P48* | Y | NA | N |
| *P49* | N | Alpha | Y |
| *P52* | N | Alpha | Y |
| *P53* | Y | NA | N |
| *P54* | Y | Alpha | N |
| *P55* | N | Alpha | Y |
| *P57* | Y | Alpha | N |
| *P58* | NA | NA | N |
| *P6_1* | N | Delta | Y |
| *P6_2* | N | Delta | Y |
| *R7_1* | N | Beta | Y |
| *R7_2* | N | Beta | Y |
| *S7_1* | N | Beta | Y |
| *S7_2* | N | Beta | Y |
| *S8_1* | N | Beta | Y |

Supplementary Table 1 : Metadata file

| | Gene | g+t+ | g+t- | g-t+ | g-t- | fisher_p | fisher_q | empirical_p | fq*ep |
|---|---|---|---|---|---|---|---|---|---|
| US-SP131-1 | PCL53_H1 | 15 | 1 | 0 | 82 | 8.76E-17 | 5.6E-12 | 4.98E-03 | 2.8E-14 |
| | PCL54_H1 | 15 | 1 | 0 | 82 | 8.76E-17 | 5.6E-12 | 4.98E-03 | 2.8E-14 |
| | PCL52_H1 | 14 | 1 | 1 | 82 | 6.82E-15 | 4.4E-10 | 4.98E-03 | 2.2E-12 |
| | LAFNNIIH_00972 | 12 | 1 | 3 | 82 | 7.05E-12 | 4.5E-07 | 4.98E-03 | 2.3E-09 |
| US-D-1 | PCL52_H1 | 13 | 2 | 0 | 83 | 1.95E-14 | 1.3E-09 | 4.98E-03 | 6.2E-12 |
| | PCL53_H1 | 13 | 3 | 0 | 82 | 1.04E-13 | 6.7E-09 | 4.98E-03 | 3.3E-11 |
| | PCL54_H1 | 13 | 3 | 0 | 82 | 1.04E-13 | 6.7E-09 | 4.98E-03 | 3.3E-11 |
| | LAFNNIIH_00972 | 11 | 2 | 2 | 83 | 5.20E-11 | 3.3E-06 | 4.98E-03 | 1.7E-08 |
| RU-RT21-19 | PCL53_H1 | 13 | 3 | 1 | 81 | 1.41E-12 | 9.1E-08 | 4.98E-03 | 4.5E-10 |
| | PCL54_H1 | 13 | 3 | 1 | 81 | 1.41E-12 | 9.1E-08 | 4.98E-03 | 4.5E-10 |
| | PCL52_H1 | 12 | 3 | 2 | 81 | 4.77E-11 | 3.1E-06 | 4.98E-03 | 1.5E-08 |
| | LAFNNIIH_00972 | 10 | 3 | 4 | 81 | 1.80E-08 | 1.2E-03 | 4.98E-03 | 5.8E-06 |
| RU-HA-1 | PCL53_H1 | 11 | 5 | 0 | 82 | 3.90E-11 | 2.5E-06 | 4.98E-03 | 1.2E-08 |
| | PCL54_H1 | 11 | 5 | 0 | 82 | 3.90E-11 | 2.5E-06 | 4.98E-03 | 1.2E-08 |
| | PCL52_H1 | 10 | 5 | 1 | 82 | 2.24E-09 | 1.4E-04 | 4.98E-03 | 7.2E-07 |
| | LAFNNIIH_00972 | 9 | 4 | 2 | 83 | 2.30E-08 | 1.5E-03 | 4.98E-03 | 7.4E-06 |
| RU-C20-1 | PCL52_H1 | 13 | 2 | 0 | 83 | 1.95E-14 | 1.3E-09 | 4.98E-03 | 6.2E-12 |
| | PCL53_H1 | 13 | 3 | 0 | 82 | 1.04E-13 | 6.7E-09 | 4.98E-03 | 3.3E-11 |
| | PCL54_H1 | 13 | 3 | 0 | 82 | 1.04E-13 | 6.7E-09 | 4.98E-03 | 3.3E-11 |
| | LAFNNIIH_00972 | 10 | 3 | 3 | 82 | 5.31E-09 | 3.4E-04 | 4.98E-03 | 1.7E-06 |
| RU-BOL1-1 | PCL53_H1 | 13 | 3 | 0 | 82 | 1.04E-13 | 6.7E-09 | 4.98E-03 | 3.3E-11 |
| | PCL54_H1 | 13 | 3 | 0 | 82 | 1.04E-13 | 6.7E-09 | 4.98E-03 | 3.3E-11 |
| | PCL52_H1 | 12 | 3 | 1 | 82 | 7.05E-12 | 4.5E-07 | 4.98E-03 | 2.3E-09 |
| | LAFNNIIH_00972 | 11 | 2 | 2 | 83 | 5.20E-11 | 3.3E-06 | 4.98E-03 | 1.7E-08 |
| HU-HO-2 | PCL53_H1 | 11 | 5 | 0 | 82 | 3.90E-11 | 2.5E-06 | 4.98E-03 | 1.2E-08 |
| | PCL54_H1 | 11 | 5 | 0 | 82 | 3.90E-11 | 2.5E-06 | 4.98E-03 | 1.2E-08 |
| | PCL52_H1 | 10 | 5 | 1 | 82 | 2.24E-09 | 1.4E-04 | 4.98E-03 | 7.2E-07 |
| | LAFNNIIH_00972 | 8 | 5 | 3 | 82 | 1.16E-06 | 7.4E-02 | 4.98E-03 | 3.7E-04 |
| FI-SKW2-1 | PCL53_H1 | 12 | 4 | 0 | 82 | 2.24E-12 | 1.4E-07 | 4.98E-03 | 7.2E-10 |
| | PCL54_H1 | 12 | 4 | 0 | 82 | 2.24E-12 | 1.4E-07 | 4.98E-03 | 7.2E-10 |
| | PCL52_H1 | 11 | 4 | 1 | 82 | 1.40E-10 | 9.0E-06 | 4.98E-03 | 4.5E-08 |
| | LAFNNIIH_00972 | 9 | 4 | 3 | 82 | 8.82E-08 | 5.7E-03 | 4.98E-03 | 2.8E-05 |
| BE-WH1-2 | PCL53_H1 | 10 | 6 | 0 | 82 | 5.72E-10 | 3.7E-05 | 4.98E-03 | 1.8E-07 |
| | PCL54_H1 | 10 | 6 | 0 | 82 | 5.72E-10 | 3.7E-05 | 4.98E-03 | 1.8E-07 |
| | PCL52_H1 | 9 | 6 | 1 | 82 | 2.99E-08 | 1.9E-03 | 4.98E-03 | 9.6E-06 |
| | LAFNNIIH_00972 | 8 | 5 | 2 | 83 | 3.32E-07 | 2.1E-02 | 4.98E-03 | 1.1E-04 |

Supplementary Table 2 : PAV results for the external abdomen attachment.

The table give the 4 first hits for each host produce by scoary. g+t+: Number of isolates that have the gene (g+) and have the trait (t+), g+t-: number of isolates that have the gene (g+) and do not have the trait (t-), g-t+: number of isolates that do not have the gene (g-) and have the trait (t+) and g-t-: number of isolates that do not have the gene (g-) and do not have the trait (t-). *fisher_p, fisher_q*: corrected and uncorrected *p*-value of Fisher's test. *empirical_p*: *p*-value of the post-hoc permutation test for the "best" gene. *fq*ep:* product of *fisher_q* and *empirical_p*.

| | PCL53 - | | | PCL53 + | | |
|---|---|---|---|---|---|---|
| | **C1** | **P20** | **C19** | **P38** | **P54** | **P55** |
| Host infected | 0 | 0 | 0 | **79.2%** | **77,9%** | **79.6%** |
| Host resistant | **100%** | **100%** | **100%** | 20.8% | 22.1% | 20.4% |

Supplementary Table 3 :
Three isolates exhibiting PCL53 (PCL53+) and 3 isolates not having it (PCL53-) were tested against 525 host clones for external abdomen attachment. The number in the table corresponds to the percentage of hosts resistant or susceptible at this site for each isolate.
